# *Chlamydomonas* State Transition Is Quiet Around Pyrenoid and Independent from Thylakoid Deformation

**DOI:** 10.1101/2021.10.27.465227

**Authors:** XianJun Zhang, Yuki Fujita, Naoya Kaneda, Ryutaro Tokutsu, Shen Ye, Jun Minagawa, Yutaka Shibata

## Abstract

Photosynthetic organisms have developed a rapid regulation mechanism called state transition (ST) to rapidly adjust the excitation balance between two photosystems by light-harvesting complex II (LHCII) movement. Though many researchers have assumed coupling of the ultrastructural dynamics of the thylakoid membrane to the ST mechanism, how ST is related to the ultrastructural dynamic of the thylakoid in *Chlamydomonas* remains elusive. To clarify the above-mentioned relation, here we used two specialized microscope techniques, observation via the excitation-spectral microscope (ESM) developed recently by us and the super-resolution imaging based on structured illumination microscopy (SIM). The ESM observation revealed a highly reversible rearrangement of LHCII-related fluorescence. More importantly, it clarified lower ST activity in the region surrounding the pyrenoid, which is the specific subcellular compartment associated with the carbon-fixation reaction. On the other hand, the SIM observation resolved partially irreversible fine thylakoid transformations induced by the ST-inducing illumination. Fine irreversible thylakoid transformation was also observed for the Stt7-kinase-lacking mutant. This result, together with the nearly equal structural changes in the less active ST regions around the pyrenoid, suggested the independence of the observed fine structural changes from the LHCII phosphorylation.

## Introduction

To cope with fluctuating illumination conditions, including light intensity, duration, and spectrum, that raise the risk of photodamage,^1–5^ photosynthetic organisms have developed various rapid energy-regulation mechanisms.^6–14^ Among such regulation mechanisms, state transition (ST) maintains the excitation balance between photosystem II (PSII) and photosystem I (PSI) through light-harvesting complex II (LHCII) movement.^10, 13, 15–22^ Normally, LHCII associates with PSII as a light-harvesting antenna (State 1, ST 1). When PSII is more excited than PSI, the unbalanced excitation leads to the reduction of the plastoquinone (PQ) pool, which causes the activation of a kinase called Stt7, resulting in phosphorylation of the stromal side of LHCII.^11, 12, 23–27^ The phosphorylated LHCII (p-LHCII) detaches from PSII, migrates to the domain of PSI, and promotes the formation of PSI-LHCI-LHCII supercomplexes (State 2, ST 2).^19, 21, 28–31^

Some studies have reported that the number of grana and the number of layers per grana change reversibly in the long-term light acclimation of higher plants.^32–34^ These morphological changes in the thylakoid membrane have been considered important for the physiology of photosynthetic organisms.^35–48^ However, it is not clear how the thylakoid membrane changes in response to short-term light acclimation. The morphological changes in the thylakoid under different light environments have been characterized many times on isolated membranes in vitro via Atomic force Microscopy^32, 41, 49^ or on vitrified thylakoids using the cryo-electron tomography technique at a molecular resolution.^40, 43, 50^ Although these techniques are powerful for providing the fine three-dimensional structures of thylakoid, they cannot provide non-invasive measurements of living cells. The ST effect has been confirmed by biochemical approaches,^19, 23, 33, 39, 51^ and its relevance to the transformation of the thylakoid architecture has also been suggested based on the results of small-angle neutron scattering (SANS) in *C. reinhardtii.*^39^ However, the connection of ST with thylakoid morphological transformation has never been fully validated because of the limitation of in vivo methodology to monitor the occurrence of ST at a small spatial scale in a living cell.

To fill the gap in visualizing the two physiological events, here we attempt to reveal the roles of the elaborate ultrastructural dynamics of the thylakoid membrane in the ST in *C*. *reinhardtii* by combining two specialized optical microscope techniques: noninvasive observation using the excitation spectral microscope (ESM) developed by us^52, 53^ and super-resolution imaging using a 3D-structural illumination microscope (3D-SIM).^54, 55^ Our custom-built ESM is capable of visualizing the local excitation spectrum, which is sensitive to the local antenna size of photosystems during ST, by monitoring changes in the chlorophyll *b* (Chl-*b*) component at room temperature (RT), and providing a lateral spatial resolution of ca. 440 nm.^53^ On the other hand, the 3D-SIM has been proven to visualize the fine structure of thylakoid membranes in *C. reinhardtii.*^49, 56^ However, the real-time observation of dynamic thylakoid transformation in vivo during rapid ST induction has never been realized.

In this study, the site-dependent occurrences of ST within single cells are accurately determined using an ESM system, and the fine morphological changes in the thylakoid membrane are visualized using a 3D-SIM. We observed LHCII-related fluorescence rearrangement and irreversible thylakoid transformation during ST induction in *C. reinhardtii*. Unexpectedly, our 3D-SIM observation clarified that the thylakoid architecture in the Stt7 mutant devoid of LHCII phosphorylation activity still showed drastic ultrastructure changes that are comparable to those of the WT cells under PSII light illumination. The observation clearly showed that large-scale thylakoid membrane dynamics do not require LHCII phosphorylation, indicating the irrelevance of fine transformations of the thylakoid to the ST activity. By combining the two in situ bioimaging techniques, we discovered the physiological relationship between the ST mechanism and the thylakoid membrane dynamics in *C. reinhardtii*.

## Results

### 1 Visualization of LHCII-related fluorescence rearrangement upon STs at the single-cell level using an ESM

The occurrence of the light-driven ST effect is confirmed in steady-state fluorescence (SI Text 1 and Figs. S1, 2) and ESM measurements (SI Text 1 and Fig. S3). The results showed a feasible realization of the ST under the present induction condition. Subsequently, we visualized the LHCII-related fluorescence rearrangement upon forward (ST 1→ ST 2) and backward (ST 1→ B-ST 1) induction periods using an ESM. Our ESM enables the high-speed acquisition of noninvasive excitation spectra at all pixels of a single-cell image within a few minutes and provides quantified fluorescence-intensity measurements.^53^ Figure 1 shows the excitation spectra and reconstructed Chl images for a WT and Stt7 mutant cell exposed to repeated ST inductions in which the ST effect was deficient in the Stt7 mutant. The whole (Wh) images were reconstructed from the integrated signal over the whole wavelength range of the excitation spectra. The Chl-*b* and Chl-*a* images were reconstructed from the integrated signal over the 640–660 nm and 660–690 nm regions, respectively. The Chl-*b*/Wh maps were obtained by dividing the Chl-*b* images by the whole images. In the present microscope system, fluorescence over the 700–750 nm spectral range is detected. Careful analysis of the fluorescence lifetime in our previous study showed that the observed excitation spectrum is mainly attributed to PSII (ca. 70%), whereas the remaining ca. 30% is attributed to PSI.^53^ Due to the dominant contribution of PSII, therefore, the induction to ST 2 results in the reduction of fluorescence signal. The ST effect can be intuitively observed at the small pixel size of 200 nm on these reconstructed images. The Chl-*b*/Wh ratio was modulated by the energy-transfer efficiency from Chl-*b* to the Chls emitting fluorescence in the 700–750 nm spectral range and the fluorescence quantum yield of the emitters. WT cells displayed a drastic enhancement of the Chl signal in the base upon ST 2 induction. The fluorescence distribution returned to a profile similar to the initial shape when the state of the cell returned to ST 1 (B-ST 1). We observed a decrease and increase in the Chl-*b* ratio in a cell in the forward and backward periods on Chl-*b*/Wh maps, respectively (Fig. 1A). The averaged spectrum over the whole image also reflected the reversible changes of Chl-*b* upon ST (Fig. 1C). These images indicated shape-memory behavior in the rearrangement of fluorescence components under ST. No such fluorescence redistribution is seen in the Stt7 mutant cells; their spectral profile hardly changed (Fig. 1B, D).

**Figure 1.**
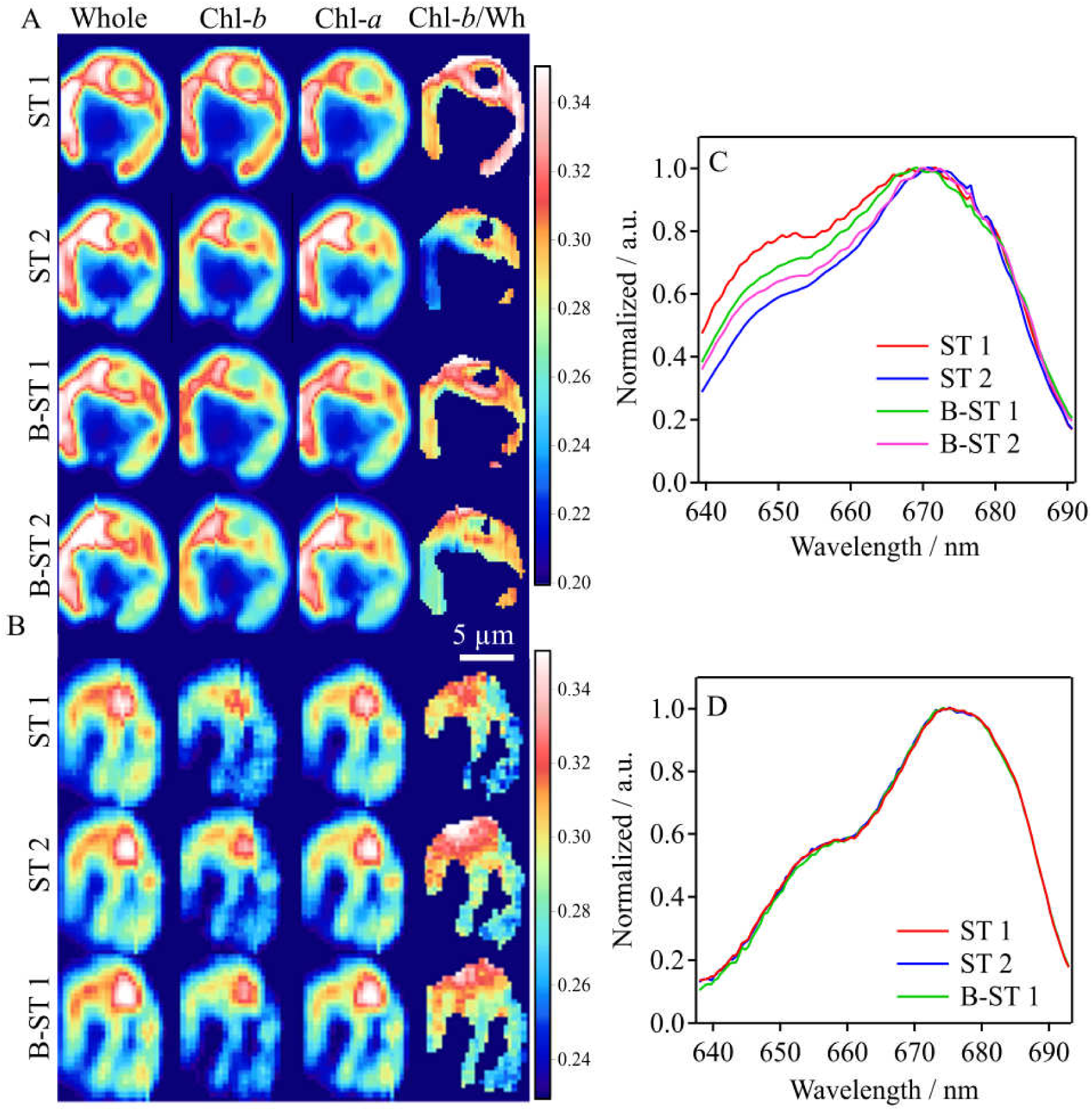
(A, B) Reconstructed images of a selected WT (A) and Stt7 cell (B) after repeated ST inductions. The “Whole” images were reconstructed from the excitation spectra over the whole wavelength regions. The “Chl-*b*” and “Chl-*a*” images were reconstructed from those integrated over the 640–655 nm and 660–690 nm spectral regions, respectively. The “Chl-*b*/Wh” images were obtained by dividing the Chl-*b* images by the Whole images. (C, D) The averaged excitation spectra over the cell for WT (C) and Stt7 (D).

To more directly show changes in the fluorescence signals in vivo during STs, we calculated the difference images between different states through subtraction. We normalized each image using a factor (whole intensity/number of pixels) so that the averaged signal intensity over the cell attains unity (Fig. 2A). It should be noted that the whole, Chl-*a*, and Chl-*b* images in the same state are divided by the same factor. This normalization eliminated the disturbance due to the signal reduction upon the transition from ST 1 to ST 2. The ST 2−ST 1 image showed drastic positive and negative signals in the region around the pyrenoid (hereafter called the basal region) and the remaining region (hereafter called lobe region), respectively. For the B-ST 1–ST 2 images, an opposite change occurred, resulting in the recovery of the initial fluorescence distribution. The difference image for B-ST 1−ST 1 was much paler, suggesting again that the fluorescence changes upon STs have good reversibility. The enhancement in the base in ST 2 was negligible in the Stt7 mutant, ensuring the relevance of the observed change to LHCII phosphorylation during STs.

**Figure 2.**
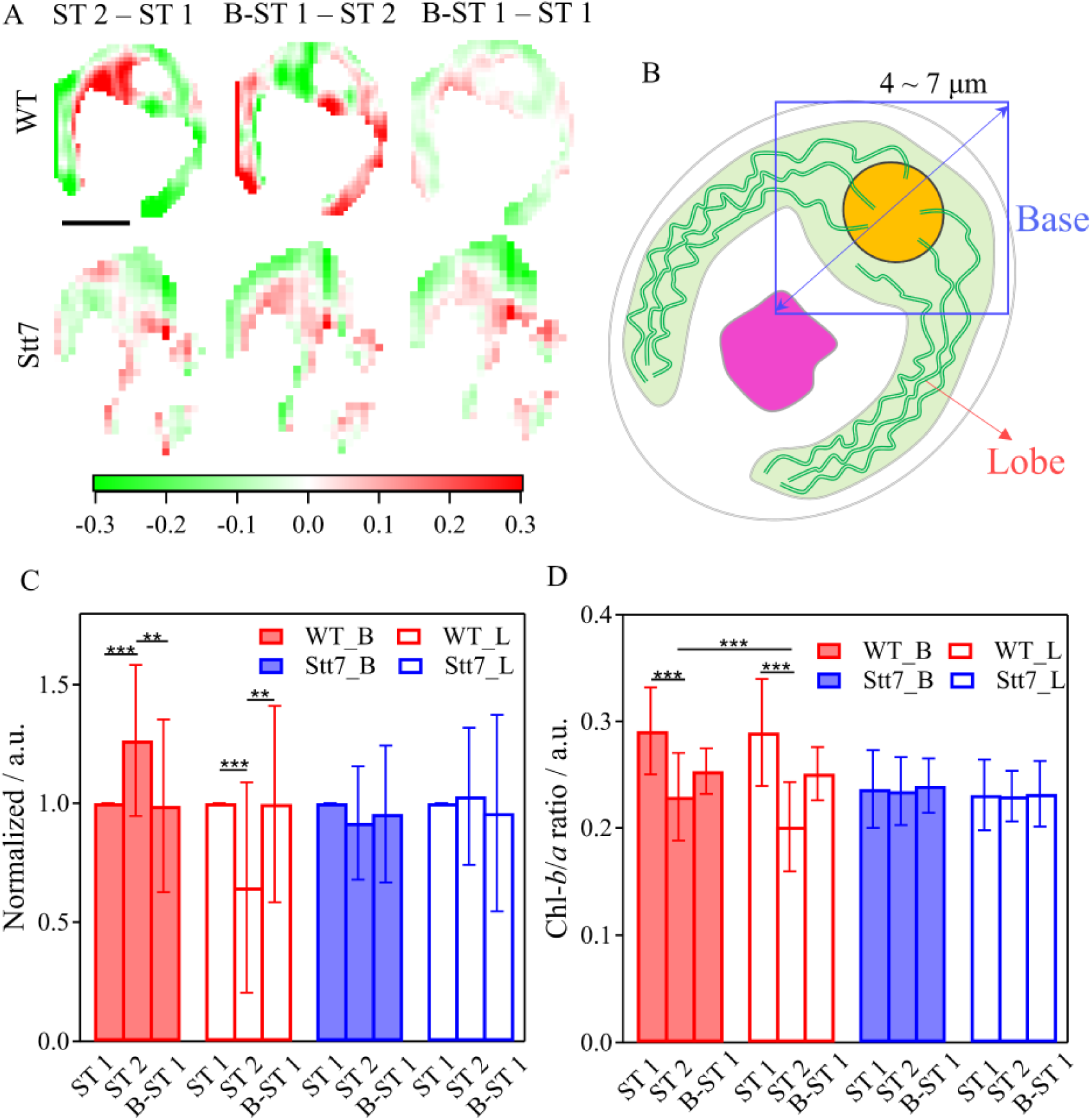
(A) Difference images of the Wh images between different states in Fig. 1 for WT and Stt7. The differences between the normalized images were calculated (see text for details). (B) Schematic view of a *C. reinhardtii* cell. The 4–7 μm square region surrounding the pyrenoid is defined as the base, and the remaining part is defined as the lobe. (C) Statistic analysis of the intensity changes of whole Chl fluorescence from WT (red, n=29) andStt7 (blue, n=19) in the base (solid) and lobe (open). (D) Chl-*b*/*a* ratio of excitation spectra of the subregions during STs summarized for WT (red, n=16) and Stt7 (blue, n=14). The Chl-*b*/*a* ratio was calculated by integrating the fluorescence intensity over 640–655 nm (Chl-*b*) and 660–690 (Chl-*a*) on the excitation spectra. The Chl-*b*/*a* ratio changes in the base and the lobe are indicated by solid and open bars, respectively. Asterisks indicate that significant differences were demonstrated using Student’s paired *t*-test (** P<0.05, *** P<0.005).

To quantify the different tendencies of the fluorescence changes between the basal and the lobe regions upon ST, in the following, we calculated the intensity changes in the two regions separately for many cells. We again used the intensity-normalized images for the calculation. As illustrated in Fig. 2B, the intensity of the base was calculated as the averaged intensity over the 4–7 μm square region surrounding the pyrenoid, and that of the lobe was calculated as the average over the remaining part. Figure 2C shows that for the WT cells, the intensity of the base is increased and decreased in the forward and backward STs, respectively, whereas opposite changes are observed in the lobe. For the Stt7 mutant cells, on the other hand, the intensities of both regions remain unchanged during the repeated ST induction treatments. The Student *t*-test analysis shows that the local intensities are significantly different between ST 1 and ST 2 and between B-ST 1 and ST 2 for WT, showing a sharp contrast to the results for Stt7.

For the cells shown in Fig. 1, we found different tendencies of decrease in the Chl-*b* ratio between the basal and lobe regions under ST induction. To quantify the changes of the Chl-*b* ratio in the base and lobe separately, we calculated the averaged Chl-*b*/*a* ratios over each region. Figure 2D shows the decrease and increase in the Chl-*b*/*a* ratio upon the forward and backward inductions for both regions. Surprisingly, the decrease in the Chl-*b*/*a* ratio upon ST 2 induction is more significant in the lobe region than in the base. This may imply ST of a different degree between the two regions, so that the layout of LHCII-related fluorescence rearrangement is influenced.

### 2 Estimation of intracellular PSII/PSI ratio

The local PSII/PSI stoichiometric ratio is of crucial importance for interpretation of the fluorescence redistribution upon ST.^18, 19, 21^ However, the imaging of the local PSII/PSI ratio has been difficult at RT.^57^ In this work, we use a PSI-specific component in the excitation spectrum to estimate the local PSII/PSI ratio at RT. As described above, the excitation spectra in the 700–750 nm range originate mainly from PSII and partly PSI.^53^ To validate the accuracy of the estimation, we prepared isolated PSII-LHCII and PSI-LHCI supercomplex mixture samples with systematically varied ratios and measured the excitation spectra of these mixtures using a steady-state instrument (Fig. S4A). The repeated measurements gave good reproducibility, and the series of spectra were globally fitted by the sum of six Gaussian functions (Fig. S4 and Table S1). The sixth component (C689) was assigned to PSI-LHCI and omitted from the fitting of the pure PSII-LHCII solution. Figure S4D shows a gradual increase in the C689 component with the increase in the PSI-LHCI ratio from 0% to 100%. We could confirm a linear relation between the enhanced amplitude of C689 and the concentration of PSI-LHCI. The PSI-specific C689 component was used to estimate the local PSII/PSI ratio in vivo.

We fitted the excitation spectrum at each pixel of an image to the sum of the six Gaussian functions with the widths and peak wavelengths fixed to the values determined in advance by the global fitting process. Then the PSI ratio map was obtained as the map of the C689 component divided by the whole image. The local PSI ratios were visualized for WT and Stt7 in Fig. S5. Figure S5D shows a statistical analysis of the PSI ratio in the base and lobe, which clarifies the tendency toward a higher PSI ratio in the base than in the lobe for both WT and Stt7, while the distribution of the PSI ratio hardly changed upon transition from ST 1 to ST 2.

### 3 In situ observation of the thylakoid membrane using a 3D-SIM

Although the ESM observation revealed changes in the Chl-*b*/Chl-*a* ratio at the local region within a chloroplast, it could not resolve the fine morphology transformations of the thylakoid, which have been considered relevant to ST. To clarify the relation of the site-dependent ST activity observed with the ESM to the fine morphology changes in the thylakoid, we observed *C. reinhardtii* cells during repeated STs using an optical super-resolution imaging technique based on the SIM technique with a lateral spatial resolution of ca. 100 nm. The 3D-SIM system used in the present study provided insight into the in situ fine morphology changes in the thylakoid ultrastructure. A 640 nm laser was used to excite mainly Chl-*b* molecules, and an emission over the 668–738 nm region was detected. The observed fluorescence originates mainly from PSII and partly from PSI (SI, Text 2– 3).

Typical images obtained with the 3D-SIM are shown in Fig. 3. Undulating stripes were frequently observed. Since the thickness of a thylakoid membrane is much smaller than the spatial resolution of ca. 100 nm, one undulating line is considered to correspond to a bundle of several thylakoid membranes. The striped patterns were more clearly observed for Stt7 (Fig. 3B, H) than for WT (Fig. 3A, G). The fluorescence distributions of WT observed with the 3D-SIM were characterized by a more disordered pattern (Fig. 3). The 3D-SIM images showed decreases of ca. 10% in the fluorescence intensities of *C. reinhardtii* cells during ST 1 to ST 2 transition, which was consistent with those observed via the ESM. No decrease in fluorescence was observed in the Stt7 mutant cells, indicating that the fluorescence reduction observed with the WT cells is ST relevant. Thus, the 3D-SIM observation enables quantitative estimation of the fluorescence intensity of live cells (Fig. S9).

**Figure 3.**
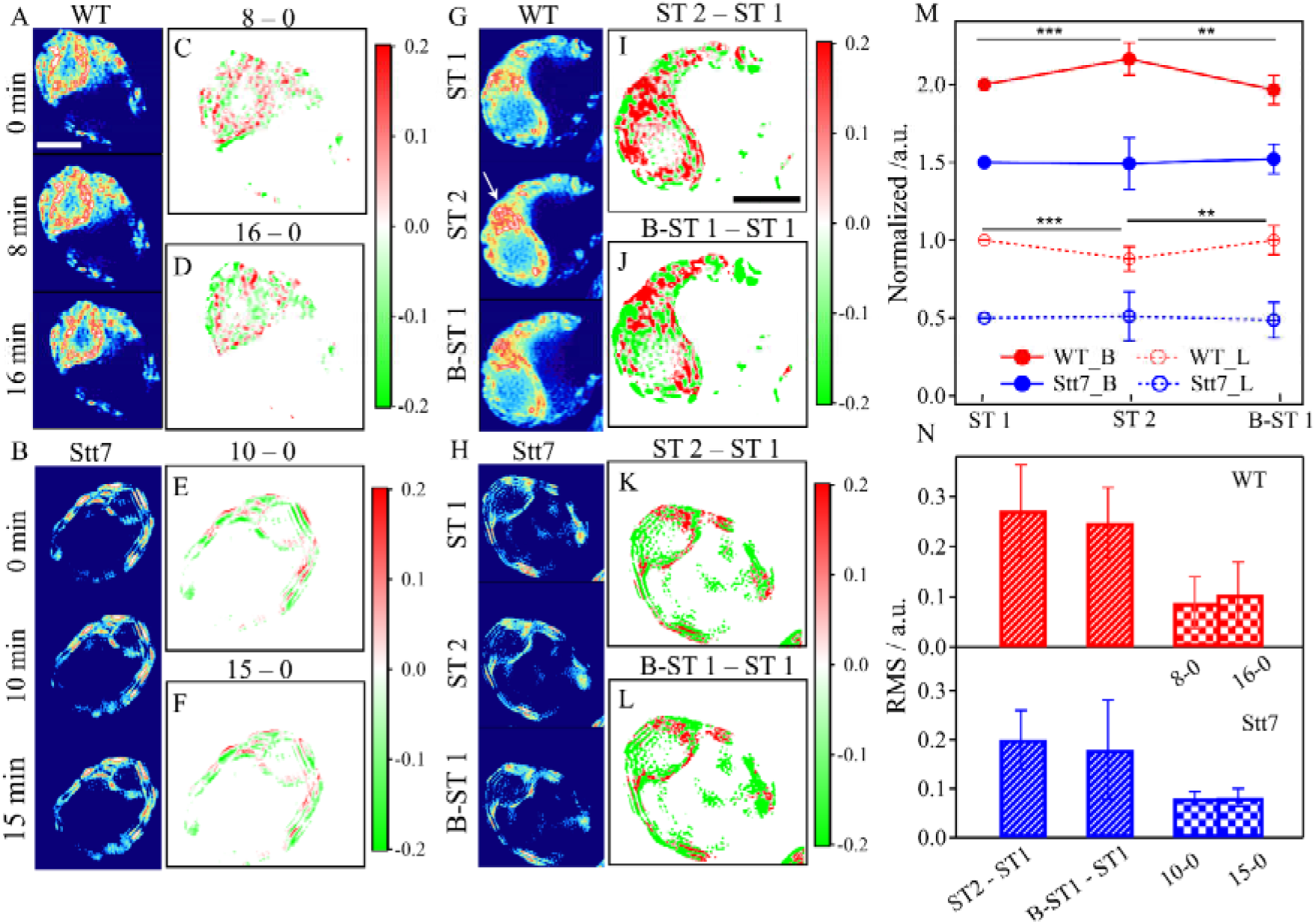
(A, B) Time course of 3D-SIM imaging for selected WT (A) and Stt7 (B) cells without ST induction. To visualize the variation in fluorescence distribution in a stationary state, we calculated the difference images at different time scales for WT (C and D) and Stt7 (E and F). (G, H) In situ observations for other WT (G) and Stt7 (H) cells in ST 1, ST 2, and B-ST 1. The difference in normalized images between the ST 2 and ST 1 (I and K, respectively) and between the B-ST 1 and ST 1 (J and L, respectively). (M) Statistical analysis of the signal intensity changes in the base (solid circles) and lobe (open circles) for the 3D-SIM images. Asterisks indicate that significant differences were demonstrated using Student’s paired *t*-test (** P<0.05, *** P<0.005); WT (red, n=13), Stt7 (blue, n=22). (N) Statistical analysis of the RMS values for the ST 2–ST 1 and B-ST 1–ST 1 images (slash bar), and the time-course images without ST induction (mosaic bar); WT (n=17), Stt7 (n=12) for the RMS calculation during ST, and WT (n=4), Stt7 (n=3) for the RMS under the spontaneous fluctuation without ST induction.

To emphasize the in situ morphological changes in the thylakoid between ST 1 and ST 2, the difference images between the states of cells were calculated as done for the ESM images, by using the normalized images. In the following section, unless otherwise stated, the fluorescence intensities of the 3D-SIM images during STs were normalized as above. Before the ST experiments were conducted, the noninvasiveness of 3D-SIM observation was confirmed by repeated measurement of the same cell without illumination to induce STs (Fig. 3A–F). We calculated the differences between images of the same cells at different times under darkness. The difference images showed random positive and negative signals over the whole chloroplast. The difference was less than 20% of the average signal and reflects the spontaneous fluctuation of the thylakoid morphology. Thus, this observation revealed a few spontaneous fluctuations of the thylakoid membrane taking place in the dark in *C. reinhardtii*. To quantify the fluctuation, we calculated the root mean square (RMS) of the difference (Fig. 3M), which showed an almost identical spontaneous fluctuation of the thylakoid morphology between WT and Stt7 (Fig. 3A–F).

Next we used the same illumination condition to induce ST as that used in the ESM measurements (Fig. 3G–L). A clear change in fluorescence redistribution was observed in WT. The cell displayed in Fig. 3G showed elaborate folding of the thylakoid membrane in the base. Milder structural changes are seen in Stt7. In the difference image of ST 2-ST 1, we found that the Chl signal shows fluorescence enhancement in the base at ST 2 for WT, which is consistent with the ESM observation (Fig. 2C). Figure 3M shows the intensity changes during ST for the base and lobe separately. The intensity in the base was enhanced and decreased in the forward and backward STs for WT. In contrast, there was no significant change for Stt7. Interestingly, the B-ST 1-ST 1 images also showed a significant change in signals throughout the whole cell. Thus, the improved spatial resolution of the 3D-SIM clearly demonstrated the irreversible nature of the fine morphological changes of the thylakoid. Such structural irreversibility could not be detected by ESM observation, whereas roughly reversible recovery of the LHCII-related fluorescence redistribution upon the backward transition to ST1 was observed.

Unexpectedly, we also observed small but non-negligible morphological changes in Stt7. The morphological changes in the thylakoid of Stt7 were significantly larger than the spontaneous fluctuation, as the deeply colored difference images in Fig. 3K, L indicate. Figure 3N shows that the RMS values of the difference images of ST 2-ST 1 and B-ST 1-ST 1 were obviously higher than those calculated for the difference images of cells kept under darkness (8-0 and 16-0 for WT and 10-0 and 15-0 for Stt7), and the RMS of WT was higher than that of Stt7 under the ST inductions. These results demonstrated that although LHCII phosphorylation enhances the ultrastructural changes in the thylakoid, it plays no essential role in the large-scale changes observed via 3D-SIM. The results indicate that the ultrastructural changes in the thylakoid membrane can be induced by light illumination only, probably through photosynthesis activity. It should be noted that the intensity of the ST-inducing light is much lower than that typically required to cause NPQ. Thus, the low light irradiation is sufficient to induce the observed thylakoid structural change.

### 4 Time-course imaging for the irreversible thylakoid via 3D-SIM

In order not to omit structural details of the thylakoid during STs, we took a 3D scan of live cells. Figure 4 shows a time course of 3D-SIM images showing the buildup of the ST-induced thylakoid deformation. These images of WT and Stt7 were selected from the complete set of 3D scans shown in Figs. S10–13. The WT cell again showed enhancement of the fluorescence signal around the pyrenoid by PSII light illumination. This enhancement was amplified with prolonged illumination. Predictably, the enhanced signals diminished to their initial intensity during the 15-minute re-induction to ST 1 (Fig. 4A–C). We did not find any significant fluorescence changes in the base for Stt7 (Fig. 4E–G). On the other hand, the zoomed-in view of the region surrounding the pyrenoid of this WT cell, shown in Fig. 4D, revealed a structure that may imply thylakoid protrusions into the pyrenoid. The protruded structure was observed only in the image of 10-minute ST 2 induction and was not observed at the other focal planes. More obviously, the Stt7 mutant showed some morphological features that clearly contrast with those of the WT cell. The thylakoid membrane in Stt7 was packed into the striped structure, and morphological changes in the thylakoid were milder, probably due to the absence of LHCII phosphorylation. Nevertheless, we observed that the Chl fluorescence was enhanced and diminished in the local lobe region during the repeated ST inductions in Stt7, accompanied by an irreversible structural deformation (white arrow, Fig. 4F). In zoomed-in view of Stt7, although the layout of the whole thylakoid membrane did not show wide changes upon ST induction, ultrastructural changes of the local thylakoid were still observed (Fig. 4H). These time-course images show the details of the fine ultrastructural change of the thylakoid membrane during STs, revealing its irreversible nature.

**Figure 4.**
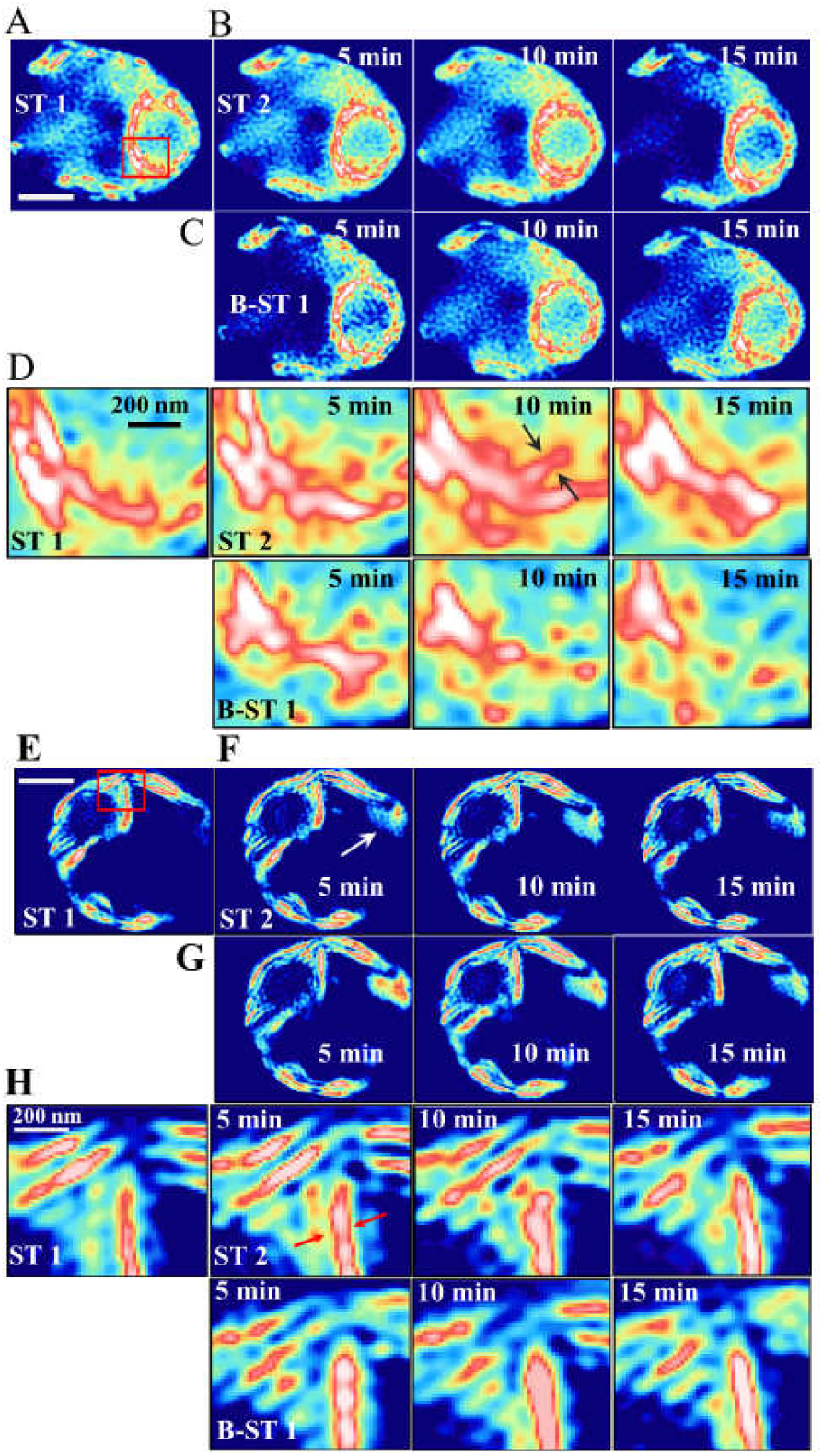
Time course of the 3D-SIM image for WT (A–D) and Stt7 (E–H) cells during ST 1 to ST 2 induction and backward transition to ST 1. (D,H) Enlarged views of the pyrenoid vicinity, corresponding to the region indicated by the red box in (A) and (E). The area marked by arrows (D) implies an appearance of the thylakoid protrusion into the pyrenoid. The areas indicated by arrows in (F) and (H) show some ultrastructural changes on the local thylakoid membrane.

## Discussion

For decades, the ST mechanism has been addressed in vivo or vitro.^18, 28, 32, 33, 35, 51, 58^ Many studies based on electron microscope observation have suggested crucial roles of the ultrastructural modification of the thylakoid upon ST.^32–34, 40, 43, 59^ However, there have been few reports on the in situ observation of dynamic ultrastructural changes in the thylakoid during ST for living cells. Therefore, the connection between the ST mechanism and the dynamic thylakoid transformation has not been fully validated. In this study, we attempted to connect the two physiological events at the single-cell level using two specialized microscopes, an ESM and a 3D-SIM, simultaneously.

With the lower resolution of the ESM, we found that the rearrangement of LHCII-related fluorescence upon repeated STs is approximately reversible (Fig. 2). The difference images show that these changes were expressed by the apparent basal enhancement in ST 2 due to the less effective reduction in fluorescence, directly reflecting changes in the local PSII antenna size (Figs. 1, 2). We confirmed that this phenomenon was caused by the onset of STs by measuring the Stt7 mutant showing no basal enhancement of fluorescence in ST 2 (Fig. 2C). These data suggest site-dependent ST activity in a single cell, which has never been reported before. We discuss possible mechanisms for this phenomenon below.

First, we consider two possible situations causing the observed apparent basal enhancement in ST 2: one in which the basal region tends to remain in ST 2 even under ST 1 induction, and one in which the basal region tends conversely to remain in ST 1 even under ST 2 induction. The former hypothesis seems consistent with the recent report by Mackinder et al. that the pyrenoid tubule accumulates PsaH which is considered to be involved in the docking of LHCII to PSI.^60^ If the former is the case, on the other hand, we should observe the lower Chl-*b* component on the excitation spectrum in the base in ST 1. This is obviously opposite to the present observation. We found that the Chl-*b*/*a* ratio was almost identical between the base and lobe in ST 1, whereas it decreased more in the lobe upon ST 2 induction (Fig. 2D). If we assume a high PSII/PSI ratio around the pyrenoid, it seems possible to explain the similar Chl-*b*/*a* ratio between the base and lobe in ST1 based on the hypothesis of the ST 2-locked base region. However, the PSI-ratio map shown in Fig. S5 indicated an opposite tendency that PSII/PSI ratio is lower in the base. Therefore, we can eliminate the possibility of the former. Instead, the basal region tends to stay in ST 1 even under a ST 2–inducing condition. Thus, the basal region seems to be less ST active.

Second, we take into account the inhomogeneity in the local PSII/PSI stoichiometry. We provided a feasible method to visualize the PSII/PSI ratio in live cells based on a deconvolution analysis of the excitation spectra. The PSI ratio maps showed a tendency for the concentration of PSI to be higher in the base than in the lobe (Fig. S6). This observation leads us to conclude that the milder reduction of LHCII-related fluorescence in the base in ST 2 comes from the lower PSII content around the pyrenoid rather than as a result of the less active ST. This assumption seems to make sense because the higher PSI content around the pyrenoid may be advantageous to the function of Rubisco. In fact, the majority of cells tended to have a higher PSI content around the pyrenoid (Fig. S5), which is in line with the above interpretation. However, there were counterexamples to this explanation: we found a non-negligible number of cells showing higher PSI ratios in the lobe together with the enhanced basal fluorescence in ST 2 (Fig. S5B). This observation clearly contradicts the above interpretation. Thus, although there is a firm tendency of PSI accumulation in the basal region, a few counterexamples suggest that the basal PSI accumulation is not a likely reason for the less significant fluorescence reduction in ST 2 in the base.

Based on the above argument by elimination, here we tentatively propose a possible explanation for the less effective basal reduction of the LHCII-related fluorescence in ST 2. We deduce that the different activity or localization of the Stt7 kinase between the base and lobe is the cause of the observed site-dependent ST activity. The activity or local concentration of the Stt7 kinase is downregulated in the base, resulting in the lower ST activity. LHCII phosphorylation is more active in the lobe thanks to the higher ST activity and makes the PSII antenna size smaller in the lobe than in the base in ST 2. There might be an unknown physiological role of maintaining the region surrounding the pyrenoid in ST 1. Biochemical and physiological validations of this interpretation will be important future endeavors.

In the 3D-SIM observation, we visualized the spontaneous fluctuation in thylakoid membranes under darkness in live *C. reinhardtii* cells. The spontaneous fluctuation of the thylakoid has also been reported in a moss species, in which a clear thylakoid fluctuation on a timescale of minutes was observed.^61^ We also clarified more ordered and striped structures of the thylakoid for Stt7 than for WT. Conversely, those of WT were found to be more folded and disordered structures, reflecting the elaborate membrane organization (Fig. 3). The more compact and ordered structure of the thylakoid in the Stt7 mutant cells might reflect the absence of its structural plasticity and flexibility. These intrinsic differences in the two strains (WT and Stt7) suggested that LHCII phosphorylation plays a crucial role in modifying the thylakoid morphology in response to long-term light acclimation.

Figure 3M shows an apparent enhancement of the basal Chl fluorescence in ST 2 in response to the ST. This result is in good agreement with the ESM observation, reflecting the roughly reversible fluorescence rearrangement in the cell upon ST. However, incomplete reversibility of morphological changes in the thylakoid was clarified via the 3D-SIM with a higher spatial resolution. Figure 4 indicates that the ST-driven thylakoid deformation was partially irreversible. Due to the nearly five-times lower spatial resolution, such fine structural information could not be obtained by the ESM. The ultrastructural changes in the thylakoid under ST induction were much larger than the spontaneous fluctuation in both WT and Stt7 cells (Fig. 3N). Surprisingly, we observed intensive thylakoid ultrastructure changes in the base, where ST activity was found to be lower (Figs. 2D, 3G). More importantly, we still observed conspicuous ultrastructural changes in the thylakoid in the Stt7 mutant cells, which are devoid of LHCII phosphorylation under PSII light illumination. Though LHCII phosphorylation in WT enhanced the thylakoid membrane reorganization during ST induction, the morphology change observed in Stt7 exceeded 70% of that of WT at the RMS level (Fig. 3N). These results indicate that the large-scale thylakoid transformations are mainly induced by the photosynthetic activity itself, not by LHCII phosphorylation, and thus are irrelevant to ST in *Chlamydomonas*. In higher plants, a kinase called Stn8 has been known to be involved in the phosphorylation of PSII core part and in the deformation of the thylakoid membrane,^62^ whereas Stn7 (the counterpart of Stt7 in *C. reinhardtii*) phosphorylates LHCII. Although the function of the counterpart of Stn8, known as Stl1 in *C. reinhardtii*, has not been fully clarified yet, the remaining Stl1 in the Stt7 mutant functions to induce the thylakoid transformation that does not result in ST 2 induction.

The thylakoid transformation during ST in *C. reinhardtii* has been reported using SANS, which offers the unique ability to make small-scale observations of ultrastructural changes in the thylakoid under ST.^39^ However, SANS revealed the averaged behavior over the ensemble of many cells but could not unveil the intracellular inhomogeneity and irreversible nature of the morphology transformation of the thylakoid. SANS data showed a small but discernible Bragg peak, which was used to characterize the appressed structure of thylakoid stacking and the inner thylakoid space in vivo with nanometer precision. The study using SANS measurement showed that the changes in the structural parameters of the thylakoid membrane by ST 1 induction were smaller in Stt7 but qualitatively similar to those in WT, which is consistent with the present observation of fine thylakoid transformations. Although the study based on SANS reported the reversibility of structural changes in the thylakoid membranes of the cell-suspension ensemble during ST 2 to ST 1 induction, our 3D-SIM imaging clearly showed that the fine structure of the thylakoid membrane of a single *C. reinhardtii* cell does not have a reversible nature during repeated ST inductions. Instead, its morphology fluctuates randomly. We observed spontaneous random deformations of the thylakoid under dark conditions (Fig. 3A–F, N). A photosynthetic reaction under ST conditions seems to enhance the random deformation of the thylakoid, resulting in its lack of a reversible nature.

It has been frequently assumed that the thylakoid membrane reorganization was intimately associated with the ST mechanism. The change in thylakoid membrane stacking was easily assumed to provide a driving force for the movement of LHCII during ST.^32–33, 39^ However, the causal relationship between the physical ultrastructural changes in the thylakoid and ST remains equivocal, since it is difficult to eliminate only the ultrastructural changes of the thylakoid itself. Based on our results, the intensity of ST activity is not correlated to the degree of thylakoid membrane transformation. We did not find any evidence that the ST mechanism and the thylakoid membrane dynamics are functionally correlated. The results of ESM and 3D-SIM observation lead us to suggest that the LHCII phosphorylation is not essential for the fine thylakoid transformation during ST that was observable by the 3D-SIM, whereas it enhances the degree of ultrastructural changes of the thylakoid. Simply, LHCII phosphorylation plays only a secondary role in thylakoid transformation, making the thylakoid structure more complicated. On the other hand, the present results do not exclude the possibility that the movement of LHCII is driven by finer thylakoid membrane dynamics than those observable by the present 3D-SIM setup (Fig. 5A).

**Figure 5.**
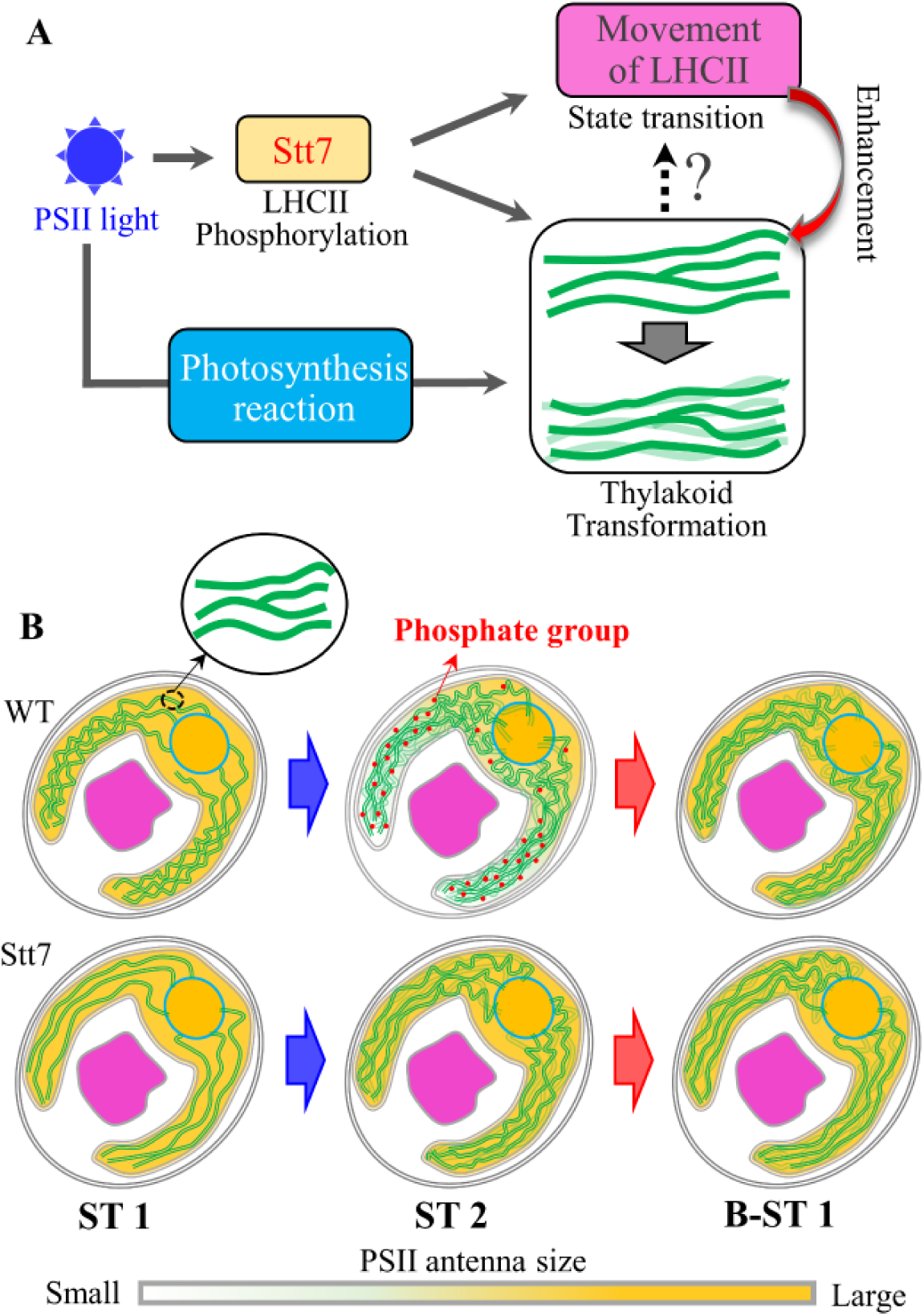
(A) The causal relation between the PSII light illumination, LHCII phosphorylation, LHCII movement, and thylakoid transformation. (B) Schematic summary of the findings of the present study. An undulating green line represents a bundle of thylakoid membranes observed using the 3D-SIM. Red dots indicate the phosphate groups attached to LHCII. The elaborate thylakoid ultrastructure does not completely recover the initial one after re-induction to B-ST 1.

Figure 5 summarizes the findings of the present study. We clarified for the first time that the fine morphology changes in the thylakoid during repeated STs are partially irreversible. We uncovered an ST-driven fluorescence rearrangement in *C. reinhardtii* for the first time. The site-dependent ST effect is visualized at the single-cell level. More importantly, for *C. reinhardtii*, LHCII phosphorylation still may be responsible for the modification of small-scale thylakoid membrane stacking, as revealed by SANS. However, our results strongly suggest that LHCII phosphorylation is not functionally associated with the large-scale ultrastructure transformation of the thylakoid in the short-term ST acclimation. This conclusion regarding a green alga seems different from those of some previous studies on higher plants that assume the necessity of LHCII phosphorylation for thylakoid membrane regulation.^33, 47^ The findings provide significant molecular bases for a deeper understanding of the two physiological events.

## Materials and Methods

### Strains and growth condition

WT *C. reinhardtii* strain 137 C and the Stt7 mutant were grown on agar plates containing Tris-acetate-phosphate (TAP) medium under low white light (20 µE·m^-2^·S^-1^). These cells were transferred to a liquid TAP medium and shaken in a rotary shaker (220–230 rpm) at 23°C with the illumination of white light (30 µE· m^-2^ · S^-1^) for three to four days. Subsequently, cells were cultivated photo-autotrophically in a liquid high-salt medium (HSM) under low white light (30 µE·m^-2^·S^-1^) for 12–24 hours before each measurement.

### Isolation of PSII-LHCII and PSI-LHCI

Thylakoid membranes from the WT were isolated as previously reported.^63^ To purify the photosynthetic supercomplexes from the isolated thylakoids, we used amphipol A8-35 as previously reported.^64^ In brief, thylakoid membranes were detergent-solubilized as described in a previous report,^63^ then immediately mixed with A8-35 at a final concentration of 1.0% and incubated on ice for 15 minutes. The A8-35-treated membranes were separated by detergent-free sucrose density gradient ultracentrifugation at 230,000×g for 16 hours.

### Light induction of STs

The ST induction light (20 µE·m^-2^·S^-1^) was supplied by a halogen tungsten lamp. Far-red light preferentially stimulating PSI and blue light preferentially exciting PSII were prepared with bandpass filters centered at 710 nm and 469 nm, respectively. The induction light was transmitted through an optical fiber bundle and directly irradiated to the samples for the steady-state and microscope measurements. We confirmed that 10 minutes of illumination is sufficient for both ST 1 and ST 2 inductions. All processes were performed at room temperature, controlled at 23 ± 1°C.

### Steady-state fluorescence measurements

The steady-state fluorescence spectra at 77 K were measured using a conventional fluorometer (F4500, Hitachi) and a home-made liquid N_2_ Dewar vessel. *C. reinhardtii* cell suspensions contained in the copper sample holder were induced to ST 1 or ST 2 at RT and then immediately immersed in the liquid N_2_ in the vessel. The optical density of the sample was adjusted to ca. 0.1 using the HSM solution.

### 3D-SIM super-resolution imaging

The super-resolution imaging for live cells was performed using a 3D-SIM microscope (N-SIM S, Nikon, Tokyo, Japan) with a water-immersion objective (×60, NA=1.27) (https://www.nikon.com/products/microscope-solutions/lineup/s-resolution/nsim/). The spatial resolutions of the microscopic system were estimated to be 123 nm and 334 nm for lateral and axial directions, respectively (for more details, see Fig. S7). The system is equipped with an advanced perfect focus system (PFS), which can automatically correct defocusing due to the intrinsic thermal drift of the optical instruments and enables the long-time in situ observation of live cells. We used a continuous wave laser at 640 nm to excite mainly Chl-*b* and partly Chl-*a*. Fluorescence over the 664–738 nm region was detected. The emission regions covered the signals of abundant PSII- and PSI-related supercomplexes. Image acquisitions were carried out using NIS-Elements AR software (Nikon, Tokyo, Japan). Each image was composed of 1024×1024 pixels, and the size of one pixel was evaluated to be 20 nm. The raw images were reconstructed to wide-field and SIM images using NIS-Elements AR software. Image analysis was also performed using Igor Pro 8 software (WaveMetrics, Tigard, OR, USA).

### Excitation spectral microscope system

A custom-built microscope system was used to rapidly obtain the intracellular excitation spectra at every pixel of the images. Our optical system has the potential to visualize the light-harvesting ability of antenna proteins at a small spatial scale and is sensitive to LHCII-related fluorescence changes on STs at RT.^52, 53^ For the details of its operation, see our previous report.^52, 53^ Briefly, an excitation light in the 600–700 nm range was generated by a photonic-crystal fiber (PCF) (SCG-800, Newport, Irvine) pumped by the Ti:S laser (MAITAI, Spectra Physics, Milpitas, USA) at 756 nm. Next, the light was dispersed by a prism to realize a wavelength-dispersed line focus at the focal plane of the objective lens. The fluorescence beyond 700 nm through a dichroic mirror (DM) (FF700-Diol-25x36, Semrock, Rochester) was injected into a polychromator (SpectraPro, 320i, Acton Optics, Acton) and detected by an electron-multiplying charge-coupled device (EMCCD) camera (Pro EM-HS-A, Princeton, Instruments, Trenton). An oil-immersion objective lens with NA=1.30 (×100, Nikon) was used in this system.

## Acknowledgements

This work was supported in part by JSPS KAKENHI Grant Numbers JP26650043, JP15H04356, JP15F15032, and JP19H03187 to Y.S. We thank Tohoku University Technical Support Center for providing the chance to use the 3D-SIM microscope.

## Supplemental Information

### Text 1

#### Confirmation of the occurrence of a light-driven ST

Figure S1 shows a typical fluorescence emission spectrum at 80 K upon 445 nm excitation, in which the PSI fluorescence (wavelength region around 715 nm) was enhanced when the PSII fluorescence (wavelength around 685 nm) was normalized. The PSI fluorescence increased thanks to the association of p-LHCII that detached from PSII (Fig. S1B). The ratio between the PSII and PSI emission bands remained constant in the Stt-7 mutant regardless of the ST induction (Fig. S1A).

The ST effect for C. *reinhardtii* cells can be also confirmed by using the excitation spectra at RT. Figure S2 shows an obvious decrease in the signal of Chl-*b* bound to LHCII (wavelength around 650 nm) in the excitation spectra, reflecting the detachment of LHCII from PSII. In contrast, the decrease in the Chl-*b* signal was not observed in the excitation spectra of the Stt-7 mutant. Additionally, there were no changes in the fluorescence emission spectra at RT for either WT or Stt-7 (Fig. S2). All experiments were performed under the same light-induction conditions (see Materials and Methods).

In the ESM measurements, fluorescence over the 700–750 nm range was detected on the excitation spectra. It was confirmed in the previous study^53^ that the fluorescence from PSII mainly (ca. 70%) contributes to the excitation spectra in the present condition. On account of the detachment of mobile LHCII from PSII during the transition from ST 1 to ST 2, the total Chl fluorescence is decreased at RT. To show the quantitative property of the ESM, we calculated the absolute intensity from individual cells. The absolute intensity decreased ca. 20% in the WT cells upon ST 1 to ST 1 transition at RT, but the intensity remained constant in the Stt-7 mutant cells (Fig. S3A). Additionally, we give a normalized excitation spectrum averaged over many cells (n=16 for WT and n=15 for Stt7). The decreased Chl-*b* component in the excitation spectrum in ST 2 could be confirmed, whereas the spectral profile of Stt-7 remained the same (Fig. S3B). The results obtained using the ESM on STs were consistent with those obtained using the steady-state instrument.

### Text 2

#### Contribution of free LHCII

Free LHCII may play an important role in the energy-regulation mechanism. We tentatively discuss whether the formation of free LHCII can contribute to the signal in 3D-SIM measurements. First, we hypothesize that the intra-chloroplast distributions of the PSII and PSI do not change upon STs based on the observation of the PSI ratio maps obtained using the ESM (Fig. S5). Next, we consider the possibility that changes in the distributions of the fluorescence observed via the 3D-SIM are contributed by the free LHCIIs, which may be more fluorescent than those in the functional state. In this case, the apparent fluorescence enhancement in the base in ST 2 reflects the accumulation of free LHCII there (Fig. 1A). If this is the case, we should have observed the enhanced relative signal of Chl-*b* bound to free LHCII in the vicinity of the pyrenoid. However, the enhancement of the Chl-*b* ratio was not observed in the Chl-*b*/Wh maps of ST 2 (Fig. 1). In fact, the Chl-*b* ratio decreased throughout the entire chloroplast upon transition from ST 1 to ST 2. Consequently, we conclude that these changes in fluorescence in the 3D-SIM images cannot be attributed solely to the free LHCII contribution.

### Text 3

#### Fluorescence composition in 3D-SIM observation

The spatial resolution of the 3D-SIM was estimated to be 334 nm and 123 nm for the axial and lateral directions, respectively (Fig. S6). The 640 nm continuous-wave laser was used mainly to excite the Chl-*b* molecules. A fluorescence emission over the 668– 738 nm range was detected. Figure S7 shows that the 3D-SIM can distinguish the fine ultrastructural changes of the intra-chloroplast thylakoid membrane that could not be distinguished by the conventional optical microscope. To determine the fluorescence contribution of each photosynthetic apparatus at RT, we measured the steady-state fluorescence emission spectra of the PSII-LHCII and PSI-LHCI supercomplexes isolated from *C. reinhardtii* (Fig. S8). The contribution of PSI-LHCI is often assumed to be negligible at RT due to due to its low FQY, but the result in Fig. S8B showed a non-negligible fluorescence signal from the isolated PSI-LHCI supercomplex. Therefore, the fluorescence emission in the 668–738 nm region was contributed mainly by PSII and partly by PSI according to 3D-SIM imaging.

**Figure S1.**
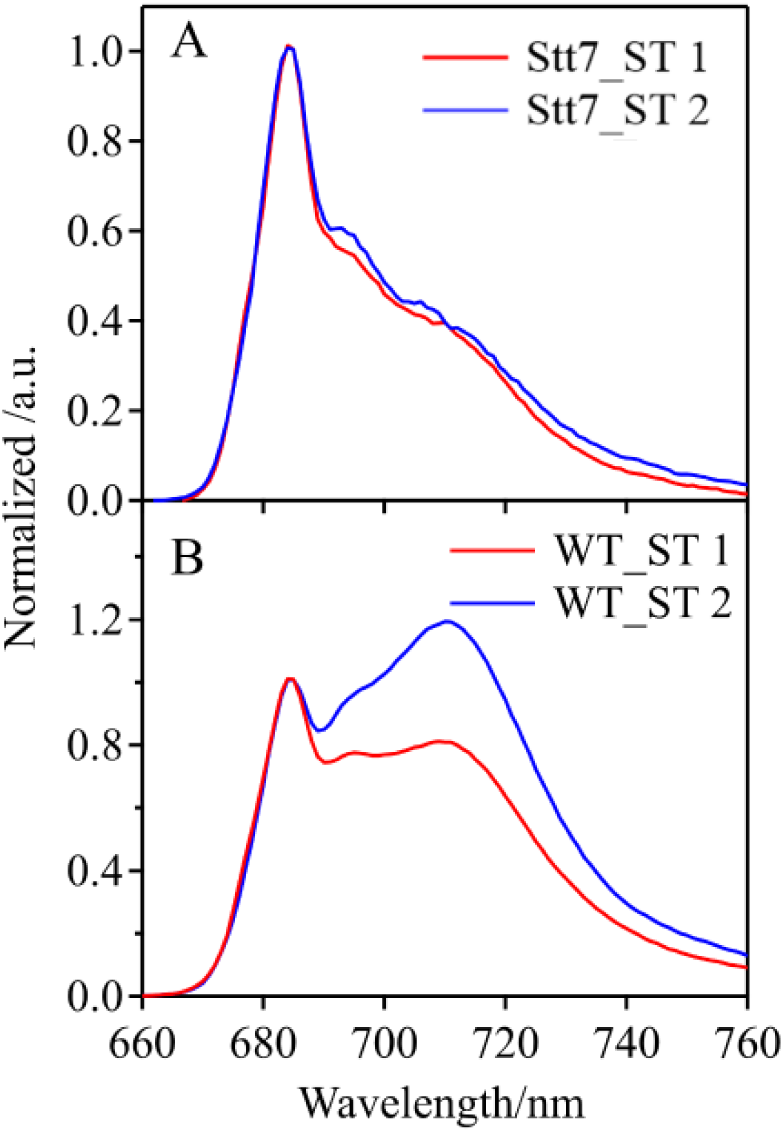
Typical fluorescence spectra, at 80 K, of Stt7 (A) and WT (B) cell suspensions excited at 445 nm. These cells were induced to ST 1 (red) and ST 2 (blue) at 23°C and immediately immersed in liquid N_2_. The cell concentration was adjusted to give ca. OD=0.1 to avoid the reabsorption effect. The peak heights in the 680 nm region on the spectra were normalized to unity. The transitions to ST 1 and ST 2 were induced by 10-minute PSI and PSII light illuminations, respectively.

**Figure S2.**
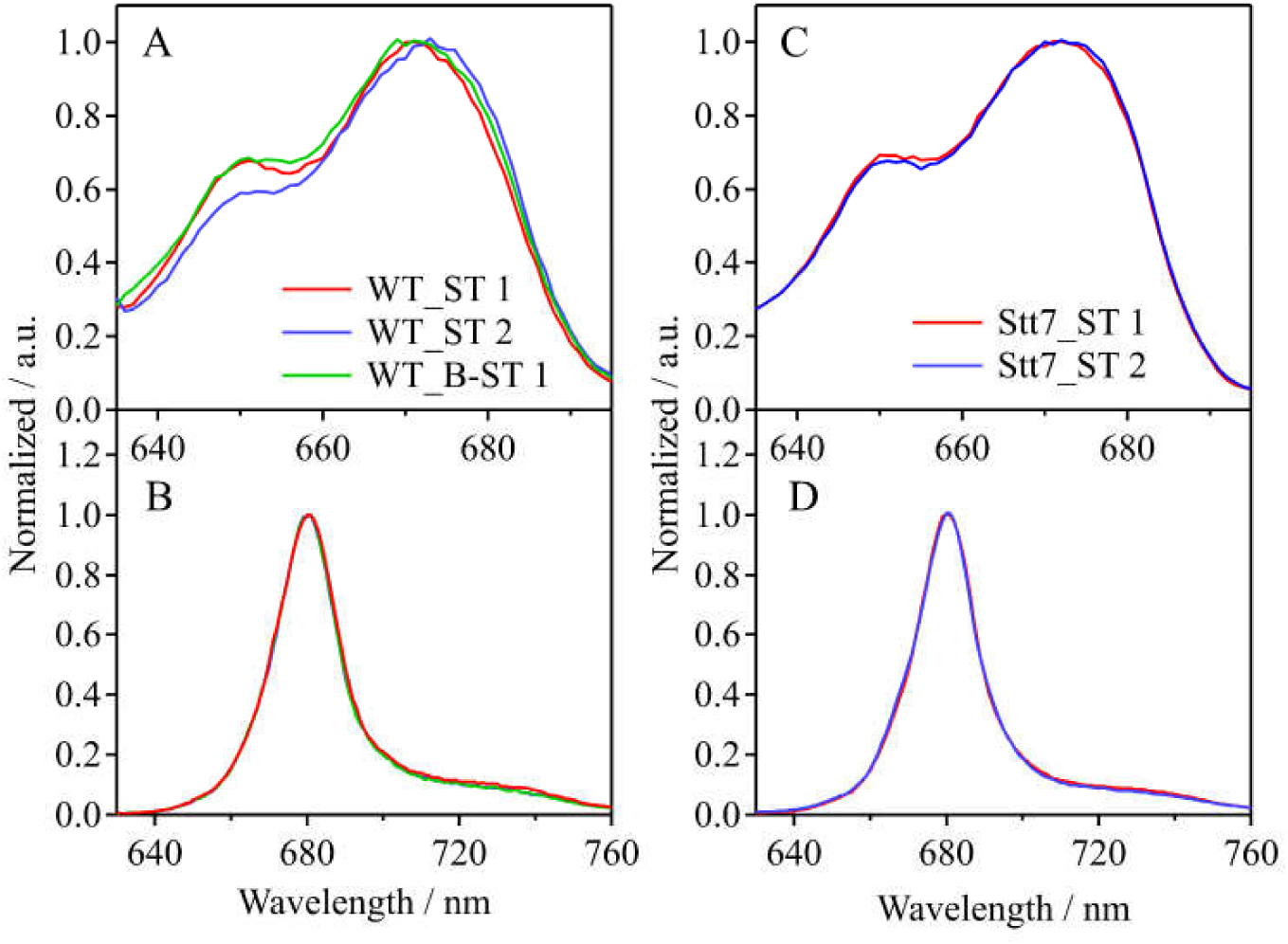
Steady-state measurements of spectra during STs at RT. Fluorescence excitation (A) and emission (B) spectra of the WT cell suspension. Fluorescence excitation (C) and emission (D) spectra of the Stt7 mutant cell suspension. An excitation wavelength of 445 nm was used to measure emission spectra. Emissions over the 700– 720 nm range were monitored for measurement of the excitation spectra. The cell concentration was adjusted to give ca. OD=0.1 to avoid the reabsorption effect in all measurements. The peak heights of the spectra were normalized to unity. The ST effect was induced by 10-minute PSI and PSII light illuminations.

**Figure S3.**
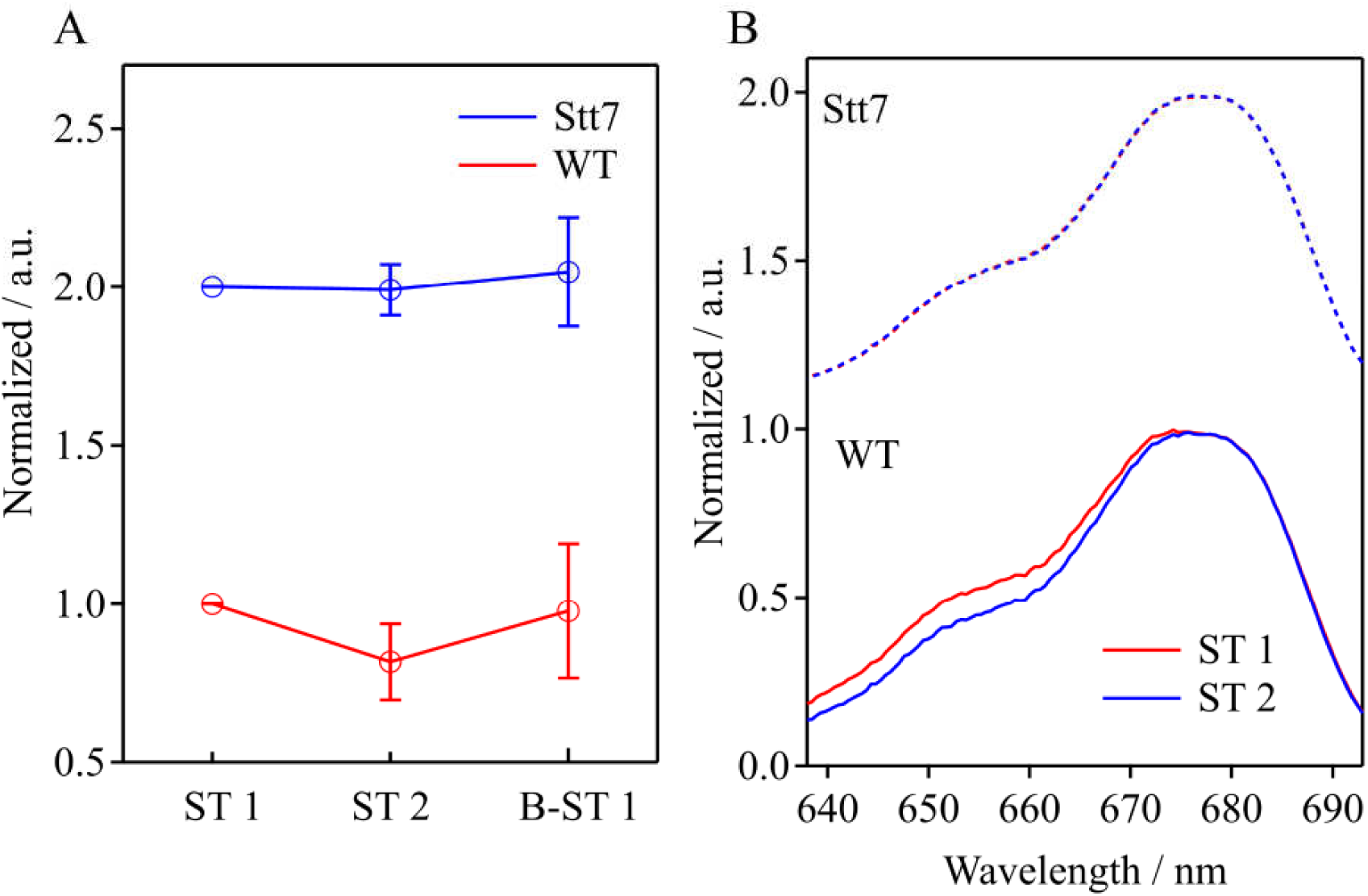
(A) Statistical analysis of the change in the fluorescence intensity during STs for the ESM images reconstructed by excitation spectra from WT (n=21) and Stt7 (n=23) upon STs. (B) Averaged excitation spectra from WT (n=16) and Stt7 (n=15) induced to ST 1 (red) and ST 2 (blue); soli lines and dotted lines indicate the WT and Stt7, respectively.

**Figure S4.**
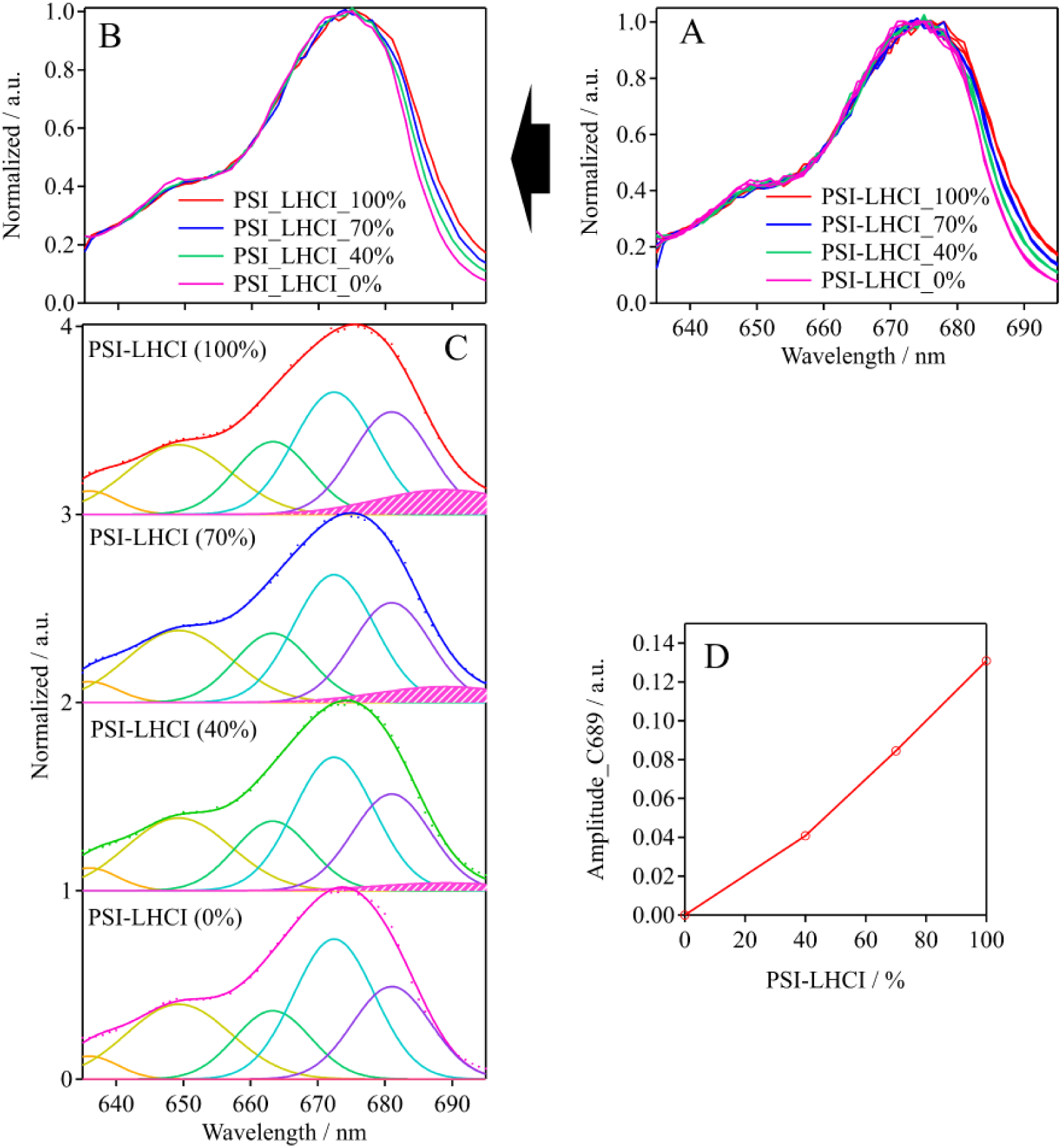
(A) Excitation spectra (n=3) of the purified PSII-LHCII and PSI-LHCI supercomplex solution samples mixed at various ratios. (B) Averages of three independent experimental results are shown. (C) The normalized excitation spectra in (A) were globally fitted to the sum of six Gaussian functions. The curves filled with magenta show the PSI-specific C689 component. (D) The amplitude of the C689 component is plotted against the ratio of the PSI-LHCI supercomplex in the mixture.

**Table S1.** Global fit analysis results for the excitation spectra of PSI-LHCI and PSII-LHCII supercomplexes. The excitation spectra of the two samples were fitted to the sum of six (for PSI-LHCI) and five (for PSII-LHCII) Gaussian functions whose widths and center wavelengths were constrained to have the same values for the spectra of both samples. The C689 component was introduced only for the PSI-LHCI spectrum and was assigned to PSI-LHCI. The values after the “±” are the 95% confidence intervals.

| Components | Peak Wavelength<br>(nm) | FWHM (nm) |
| --- | --- | --- |
| C636 | $636 \pm 1$ | $5.32 \pm 1$ |
| C649 | $648.7 \pm 1.5$ | $10.61 \pm 0.75$ |
| C664 | $663.5 \pm 0.75$ | $8.5 \pm 0.5$ |
| C672 | $672.3 \pm 0.5$ | $8.5 \pm 0.5$ |
| C680 | $680.2 \pm 1.5$ | $8.5 \pm 0.5$ |
| C689 (PSI) | $689 \pm 1$ | $14.13 \pm 0.5$ |

**Figure S5.**
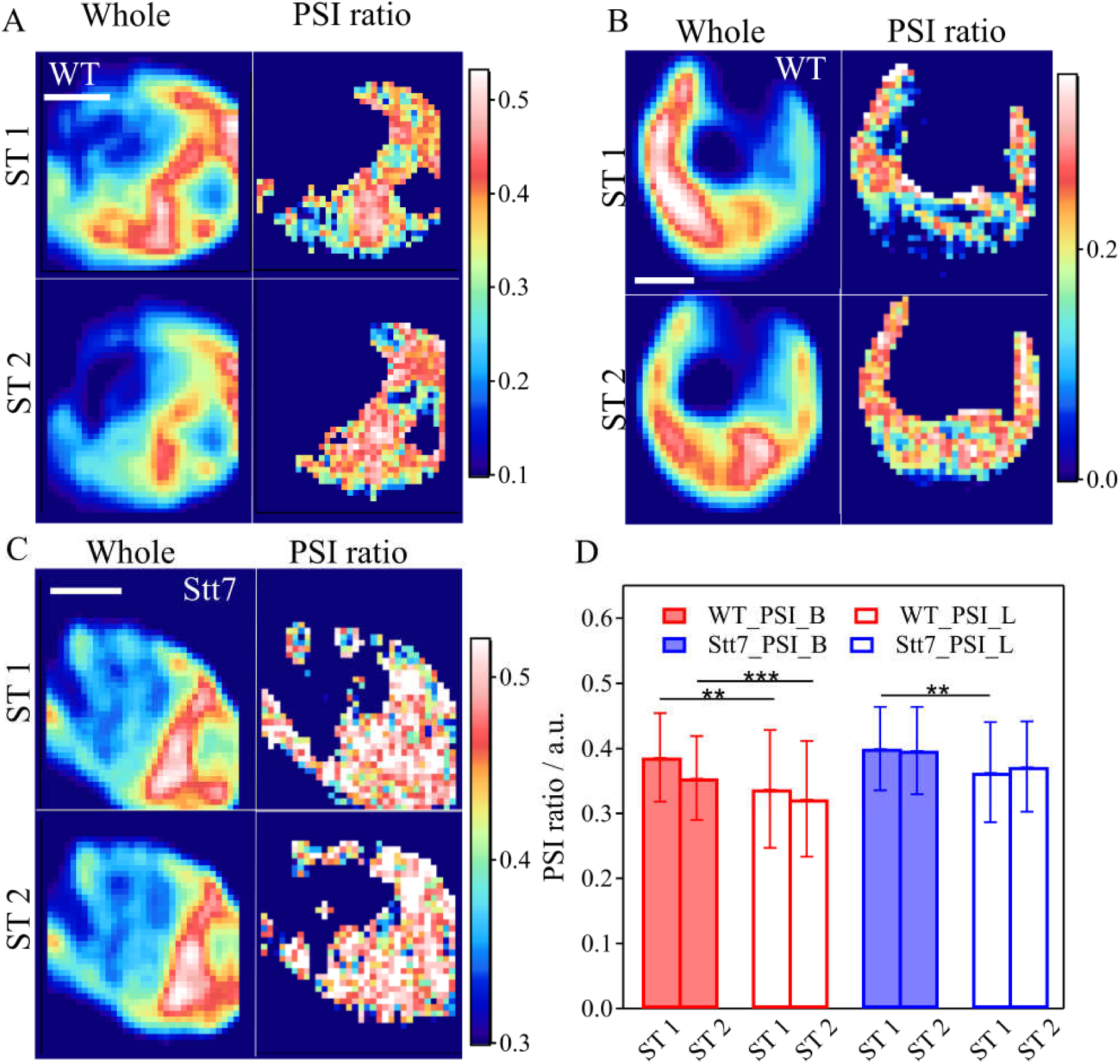
(A–C) Whole images (left) and the reconstructed PSI ratio maps (right) obtained using the PSI-specific C689 components for selected WT (A, B) and Stt-7 (C) cells. The white bar indicates 2 μm. (B) A particular WT cell whose PSI ratio is exceptionally high in the lobe and still shows the apparent basal fluorescence enhancement in ST 2. (D) Statistical analysis of the reconstructed PSI ratio maps during STs. The PSI ratios in the base and lobe regions are indicated by solid and open bars, respectively. WT (n=21), Stt-7 (n=13). ** P<0.05, *** P<0.007; the P values are obtained from Student’s paired *t*-test.

**Figure S6.**
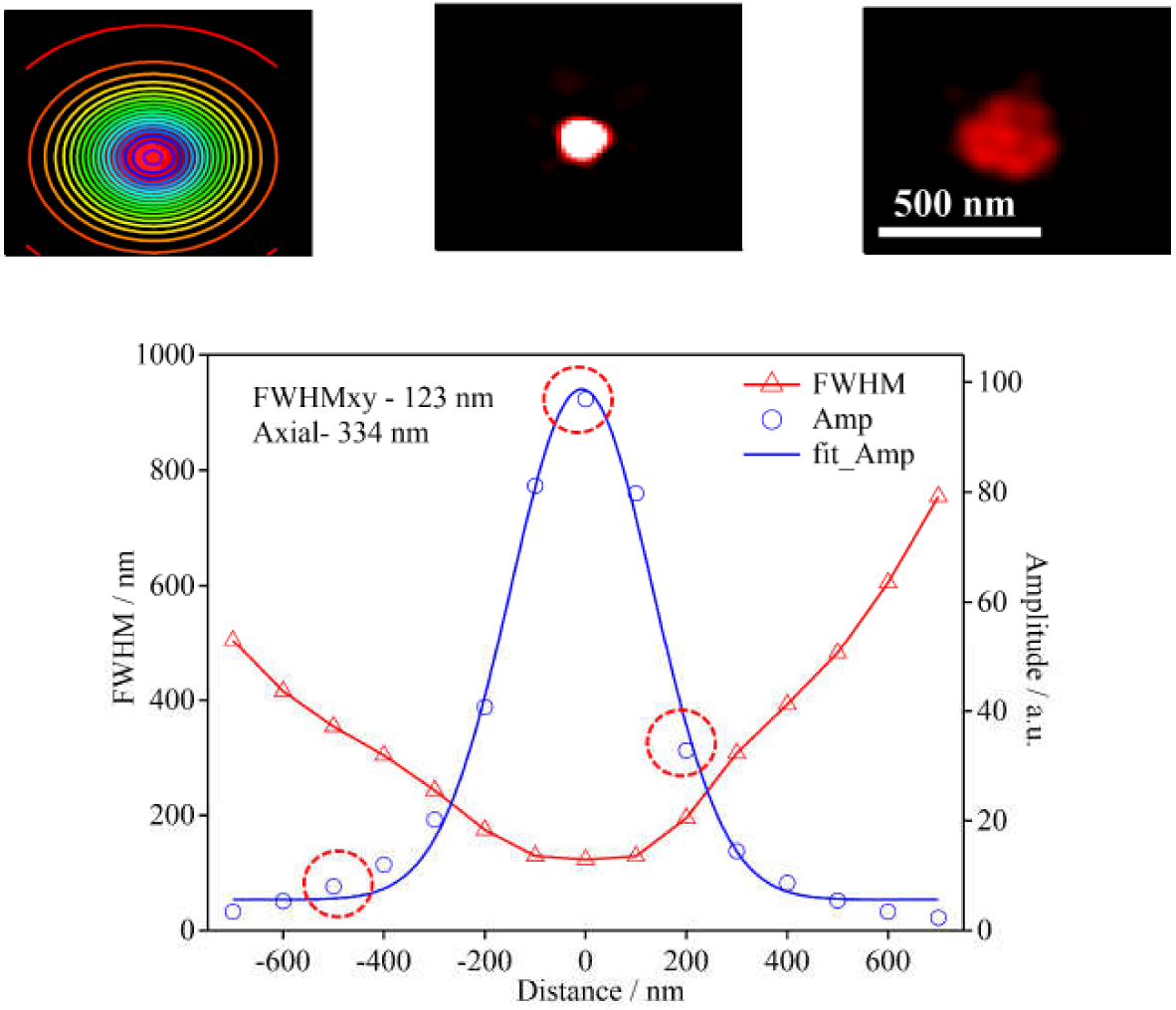
The point spread function (PSF) was obtained by measuring the fluorescence beads with a radius of 100 nm using the 3D-SIM microscope with a 100 nm axial interval. The circles and triangles show the amplitudes and FWHMs of the 2D Gaussian functions fitted to the images, respectively. The axial-shift dependence of the amplitude of the 2D Gaussian function was fitted to a Gaussian function (blue curve). The 3D-SIM microscope produced excellent spatial resolutions of 334 nm and 123 nm for the axial and lateral directions, respectively.

**Figure S7.**
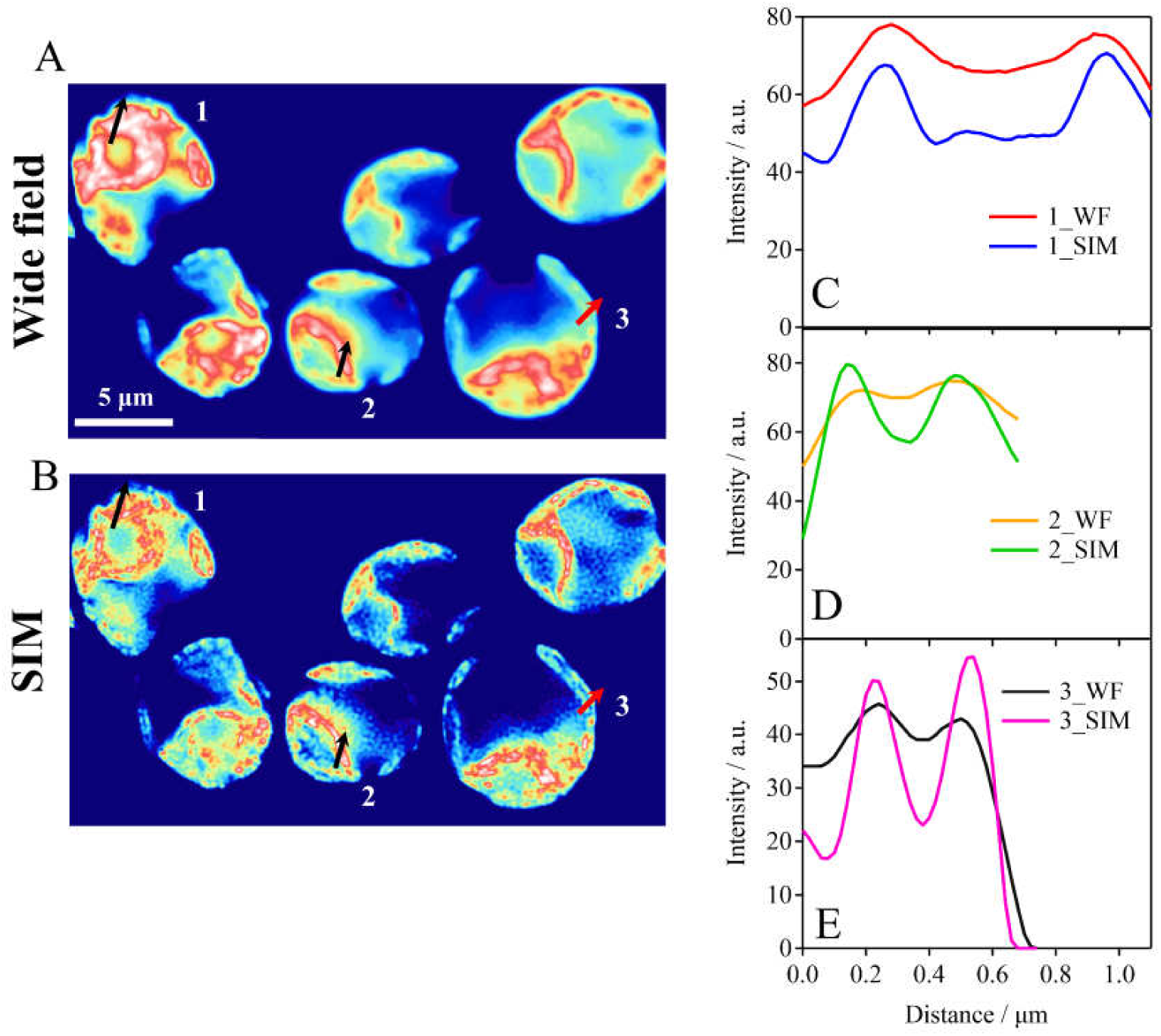
(A,B) Comparison of the reconstructed wide-field (WF) (A) and 3D-SIM images (B) of live *C. reinhardtii* cells. (C–E) Three sectional views of the fluorescence intensity along the selected region indicated by thick arrows in (A) and (B).

**Figure S8.**
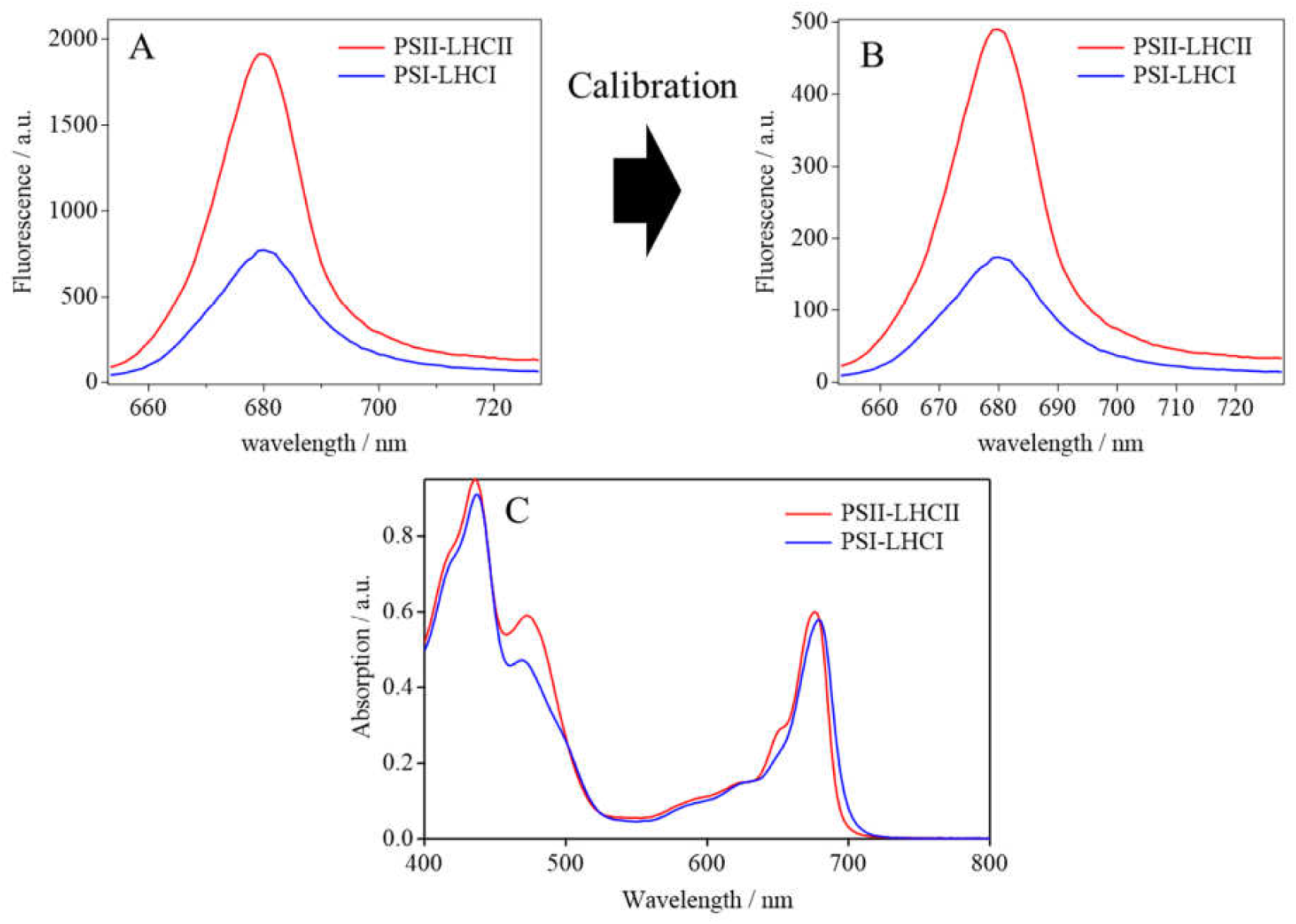
(A) Fluorescence spectra of the isolated PSII-LHCII and PSI-LHCI supercomplexes. The excitation wavelength was set to 445 nm. (B) Spectra corrected to simulate those obtained by the 640 nm excitations. To simulate the fluorescence spectra, we corrected the spectra shown in panel (A) by multiplying the factors of A640/A445 in the absorption spectra (C). The concentrations of both samples were adjusted to 1 µg and 100 µg Chl/mL using a Tritine buffer (pH 7.5, 25 mM Tritine– NaOH with 50 mM NaCl).

**Figure S9.**
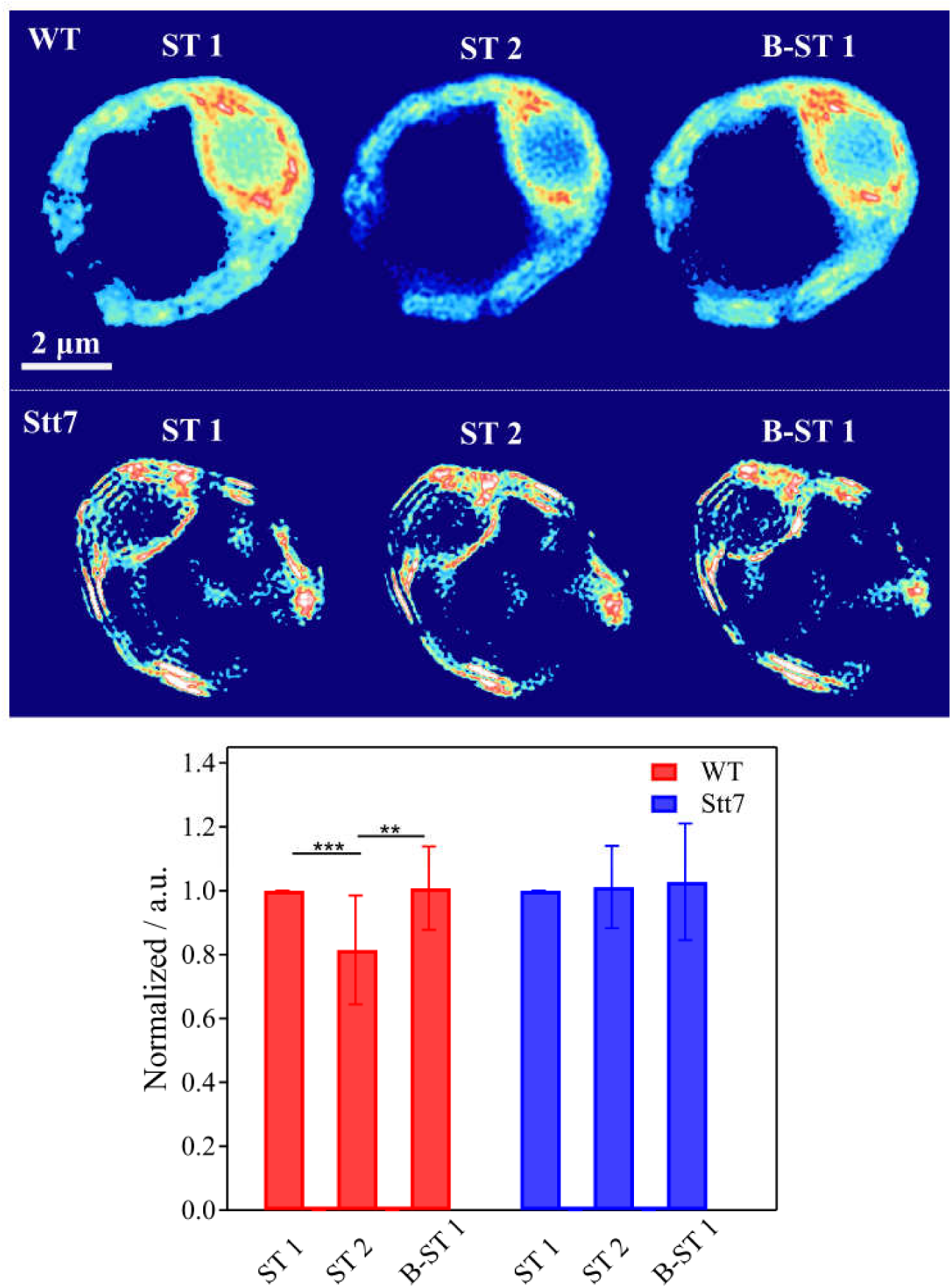
Top panel: raw 3D-SIM images of a live WT and Stt-7 mutant cell during STs. Bottom panel: average fluorescence signal intensities from the two strains (WT, n=15; Stt-7, n=12) during STs. The fluorescence intensity of cells in ST 1 was normalized to unity. The significant difference between these data was tested by using Student’s paired *t*-test: ** P<0.005, *** P<0.0005.

**Figure S10.**
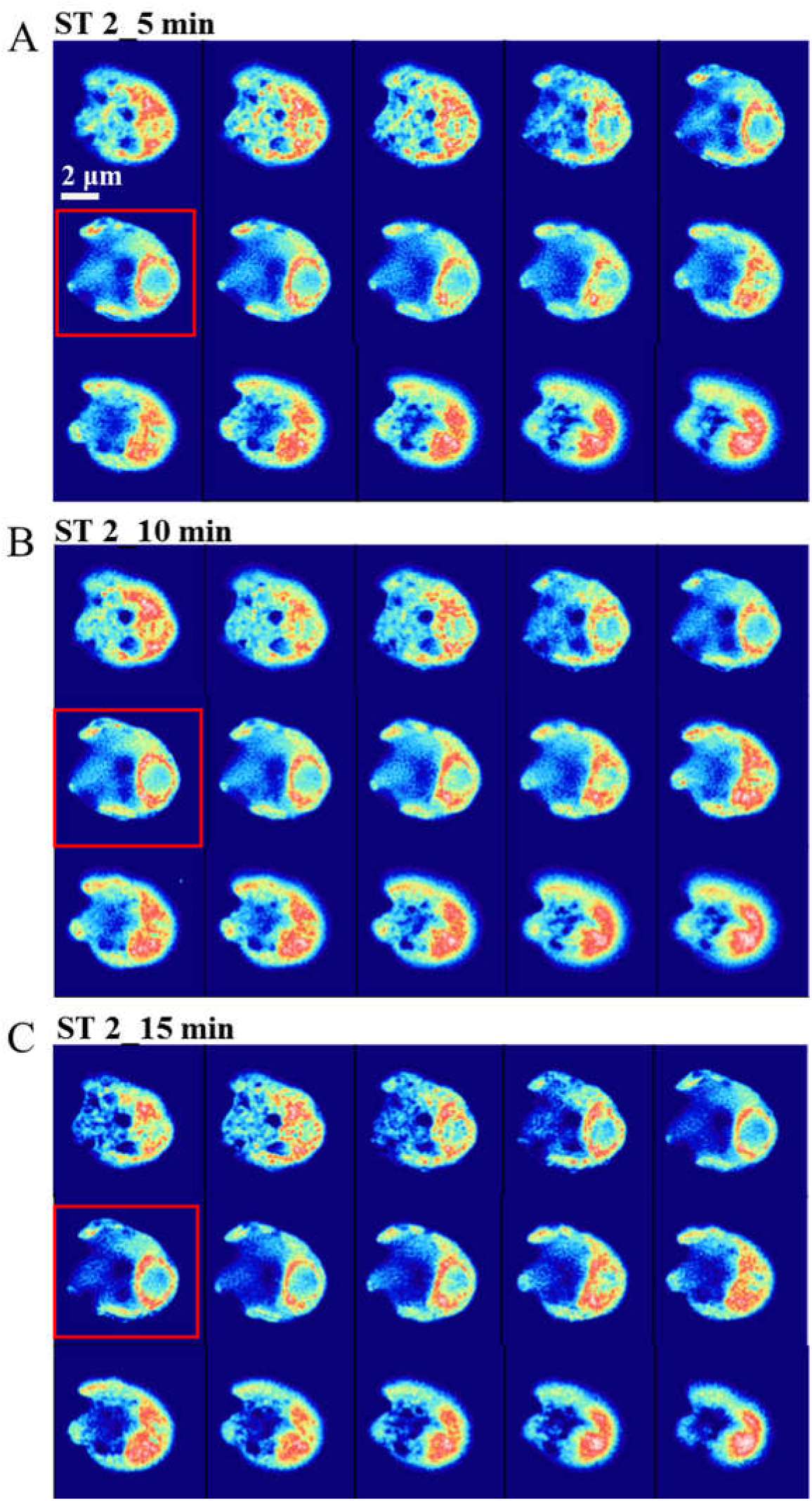
Raw 3D-scanning images of a selected WT cell upon ST 2 induction for 5 (A), 10 (B), and 15 (C) minutes. The Z interval was set to 0.25 µm. The 3D images were obtained via optical sectioning. Optical sectioning from bottom to top is shown in these images from top left to bottom right. The images indicated by red boxes were used in Fig. 4A-C.

**Figure S11.**
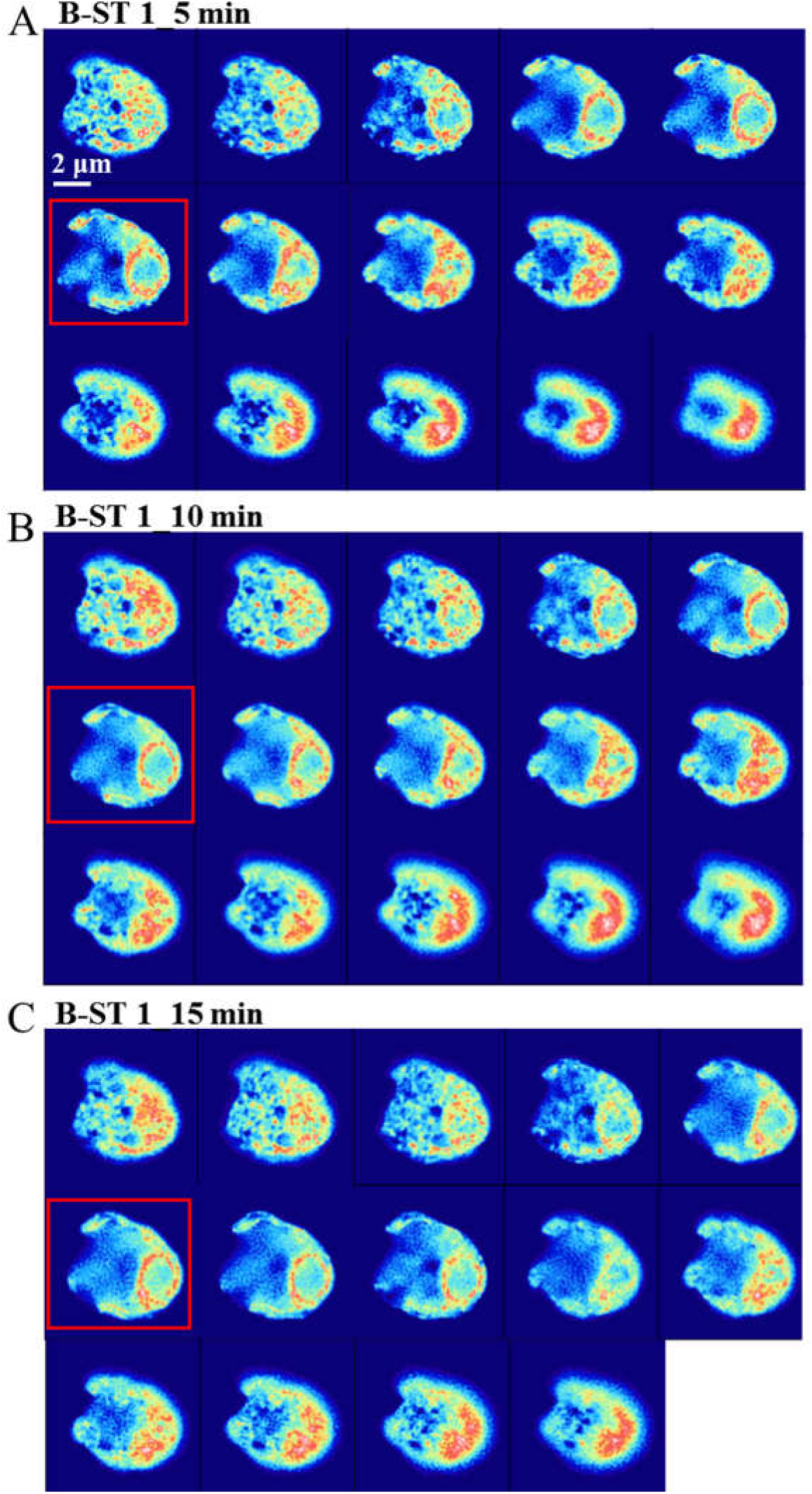
Raw 3D-scanning images of a selected WT cell upon B-ST 1 induction for 5 (A), 10 (B), and 15 (C) minutes. The Z interval was set to 0.25 µm. The 3D images were obtained via optical sectioning. Optical sectioning from bottom to top is shown in these images from top left to bottom right. The images indicated by red boxes were used in Fig. 4A-C.

**Figure S12.**
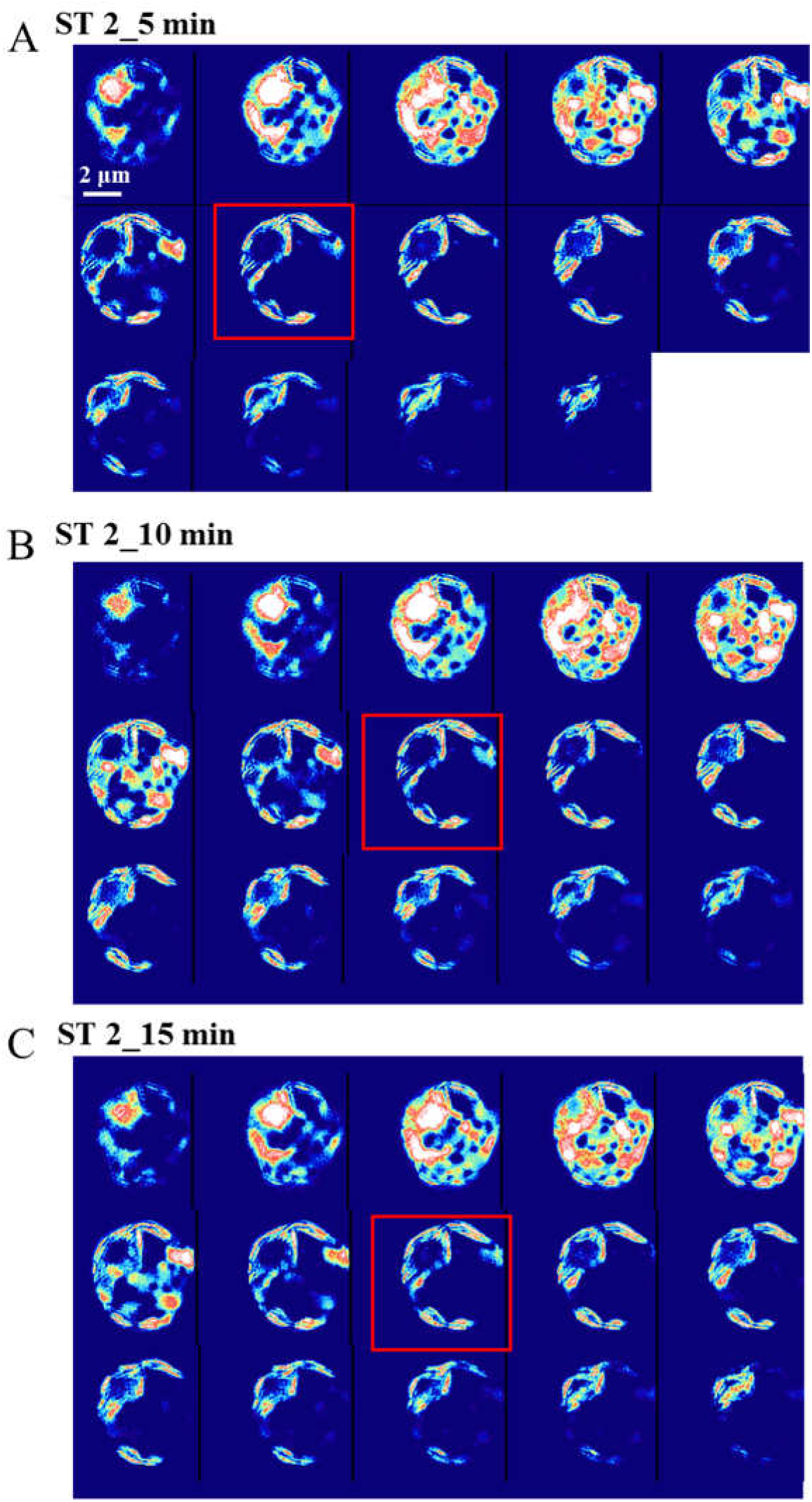
Raw 3D-scanning images of a selected Stt7 cell upon ST 2 induction for 5 (A), 10 (B), and 15 (C) minutes. The Z interval was set to 0.25 µm. The 3D images were obtained via optical sectioning. Optical sectioning from bottom to top is shown in these images from top left to bottom right. The images indicated by red boxes were used in Fig. 4E-H.

**Figure S13.**
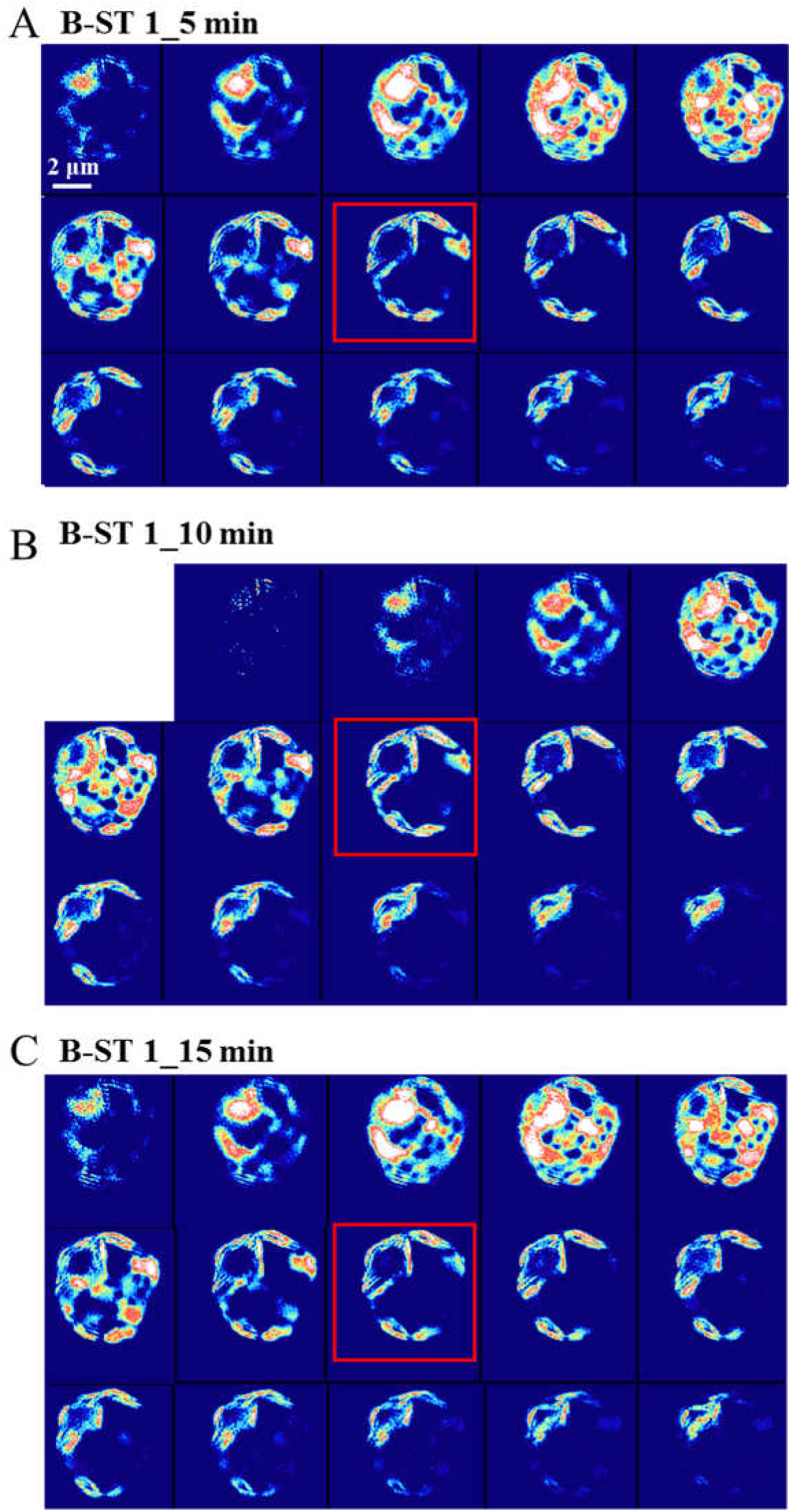
Raw 3D-scanning images of a selected Stt7 cell upon B-ST 1 induction for 5 (A), 10 (B), and 15 (C) minutes. The Z interval was set to 0.25 µm. The 3D images were obtained via optical sectioning. Optical sectioning from bottom to top is shown in these images from top left to bottom right. The images indicated by red boxes were used in Fig. 4E-H.

## Notes

### Competing Interest Statement

The authors have declared no competing interest.

